# Tripartite extended amygdala - basal ganglia CRH circuit drives arousal and avoidance behaviour

**DOI:** 10.1101/2022.03.15.484057

**Authors:** Simon Chang, Federica Fermani, Chu-Lan Lao, Lianyun Huang, Mira Jakovcevski, Rossella Di Giaimo, Miriam Gagliardi, Danusa Menegaz, Alexander Adrian Hennrich, Michael Ziller, Matthias Eder, Rüdiger Klein, Na Cai, Jan M. Deussing

**Author notes:** Corresponding Author: Max Planck Institute of Psychiatry, Molecular Neurogenetics Kraepelinstr. 2-10, 80804 Munich, Germany.

## Abstract

An adaptive stress response involves various mediators and circuits orchestrating a complex interplay of physiological, emotional and behavioural adjustments. We identified a population of corticotropin-releasing hormone (CRH) neurons in the lateral part of the interstitial nucleus of the anterior commissure (IPACL) – a subdivision of the extended amygdala, which exclusively innervate the substantia nigra (SN). Specific stimulation of this circuit elicits arousal and avoidance behaviour contingent on CRH receptor 1 (CRHR1) located at axon terminals in the SN, which originate from external globus pallidus (GPe) neurons. The neuronal activity prompting the observed behaviour is shaped by IPACL^CRH^ and GPe^CRHR1^ neurons coalescing in the SN. These results delineate a novel tripartite CRH circuit functionally connecting extended amygdala and basal ganglia nuclei to drive arousal and avoidance behaviour.

**One-Sentence Summary:** Brain centres involved in emotional and motor control are connected through a stress peptide promoting arousal and avoidance behaviour

## Main

Striving for homeostasis, the body reacts to real or perceived stressors through coordinated activation of various effector systems and brain circuits (*1*). Severe or chronic stress combined with a maladaptive stress response are well-known risk factors for neuropsychiatric disorders, such as anxiety, depression and addiction (*2, 3*). Corticotropin-releasing hormone (CRH), a central mediator of the stress response, is released in response to salient environmental stimuli activating the neuroendocrine stress axis and promoting arousal, anxiety- and fear-related behaviours (*4–10*). Moreover, the cross-talk between CRH and midbrain dopaminergic systems links the stress response to dopamine (DA) regulation (*5, 9, 11, 12*). CRH modulates the release of DA producing a positive affective state (*5, 9, 13*). However, severe stress can switch the CRH effect from appetitive to aversive (*6*). Midbrain DA neurons of the ventral tegmental area (VTA) and substantia nigra (SN) are part of the basal ganglia circuitry controlling reward function, movement initiation and coordination, which are fundamentally involved in shaping an adaptive response to stressful events (*14–16*). This response is intrinsically tied to physiological arousal, which is essential for fast adaptation including decisions on whether to approach or avoid a threat (*17*). Central application or overexpression of CRH have been demonstrated to induce arousal and behavioural activation paralleling a physiological stress response (*18–20*). However, the direct role of the CRH system and thereby governed circuits modulating the DA system in arousal and approach/avoidance behaviour are not yet understood.

Using CLARITY on brains of CRH-Cre::Ai9 reporter mice revealed an ensemble of CRH neurons at the transition between the central amygdala (CeA) and bed nucleus of the stria terminalis (BNST) defined as the interstitial nucleus of the posterior limb of the anterior commissure (IPAC), a structure previously assigned to the extended amygdala (*21*). Particularly, the lateral division of the IPAC (IPACL) contains CRH^+^ neurons in numbers comparable to the CeA and BNST (Fig. 1, A **and** B **and Movie S1 and S2**). Patch-clamp recordings and immunohistochemistry characterized them as GABAergic spiny projection neurons (*22*) with CRH^+^ neurons showing a higher excitability compared to CRH^-^ neurons (Fig 1., C **to** E **and** fig S1, A **to** I).

**Figure 1:**
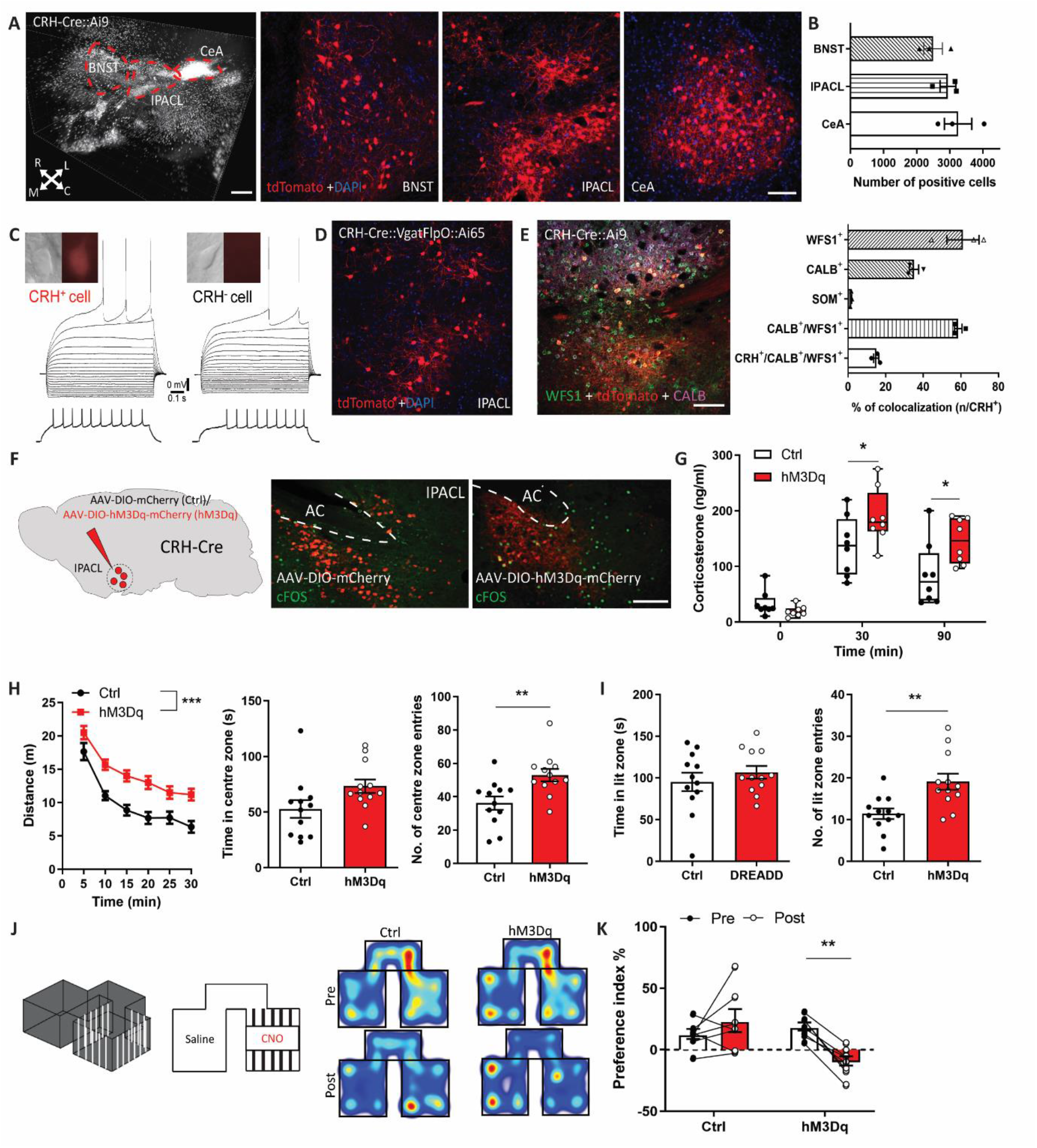
IPACL^CRH^ neurons modulate arousal and avoidance behaviour. (**A**) Snap-shot of a cleared CRH-Cre::Ai9 brain showing CRH expressing neurons throughout the extended amygdala. Representative images showing tdTomato-expressing CRH^+^ neurons in subdivisions of the extended amygdala. scale bar = 500 µm. (**B**) Quantification of CRH neurons in the CeA, IPACL and BNST. (**C**) Whole-cell patch-clamp recordings demonstrate that both CRH^+^ and CRH^-^ IPACL neurons display strong inward rectification, a depolarizing ramp potential (black arrow heads) and regular firing characteristics, indicating that they are GABAergic spiny projection neurons. (**D**) Tdtomato-expressing neurons in CRH-Cre::Vgat-FlpO::Ai65 intersectional reporter mice demonstrate that IPACL^CRH^ neurons are GABAergic. (**E**) Representative images of IPACL^CRH^ neurons co-stained with markers of GABAergic neurons and quantification of co-localization. (**F**) Scheme illustrating injection of control (AAV_8_-hSyn-DIO-mCherry) and hM3Dq-expressing (AAV_8_-hSyn-DIO-hM3Dq-mCherry) virus into the IPACL. Representative images of coronal IPACL sections, scale bar = 200 µm. (**G**) Plasma corticosterone levels at baseline, 30 and 90 min following CNO administration (repeated measures two-way ANOVA, *F*_2,42_ = 42.7, *p* < 0.0001; Bonferroni’s multiple comparison test, 30 min, *p* = 0.047; 90 min, *p* = 0.03). (**H-K**) CNO treatment induces arousal, delayed habituation and avoidance behaviour in hM3Dq-expressing animals. (**H**) Distance travelled (n = 12 repeated measures two-way ANOVA, *F*_1,22_ = 17.49, *p* = 0.0004), time spent in centre (*p* = 0.5, t = 2.1) and transitions into the centre zone (*p* = 0.005, t = 3.1) of the open field. (**I**) Time spent in lit zone (*p* = 0.4, t = 0.85) and entries into lit zone (*p* = 0.003, t = 3.3) of the dark-light box. (**J**) Schematic illustration of place preference paradigm. Heat maps illustrating the accumulated time animals stayed in respective chambers pre and post conditioning. (**K**) Calculated preference index (two-way ANOVA, Bonferroni’s multiple comparison test, *p* = 0.006, t = 3.70). Values represent mean ± SEM, * *p* < 0.05, ** *p* < 0.01, *** *p* < 0.0001.

To interrogate the function of this previously unexplored population of CRH^+^ neurons, we used Cre-dependent AAVs to express hM3Dq in IPACL^CRH^ neurons of CRH-Cre mice, which are activated upon CNO treatment (Fig. 1F **and** fig. S2, A and B). As a mediator of the stress response, CRH serves as a secretagogue triggering hypothalamic-pituitary-adrenal (HPA) axis activity and as a neuromodulator, it is implicated in the expression of stress-related behaviours. Activation of IPACL^CRH^ neurons in hM3Dq-expressing mice resulted in hyperactivation of the HPA axis indicated by increased plasma corticosterone levels 30 and 90 min after CNO administration (Fig. 1G). To determine the temporal kinetics of behavioural responses to CNO-induced IPACL^CRH^ neuron activation, mice were subjected to a 90 min open field test (OF). Mice expressing hM3Dq showed significantly elevated locomotor activity compared to control mice, peaking 15 min after CNO administration (fig. S2C). Based on this observation, we conducted different behavioural tasks assessing locomotor activity, exploratory and anxiety-related behaviour. CNO-treated mice expressing hM3Dq showed hyperactivity in the OF and the dark-light box (DaLi) (Fig. 1, H **and** I). These behavioural results, together with the increased activation of the HPA axis, suggest that activated IPACL^CRH^ neurons trigger behavioural activation and arousal. To further substantiate this hypothesis, we additionally conducted a conditioned place preference paradigm (CPP). hM3Dq-expressing mice showed clear avoidance of the CNO-paired chamber, while no difference in preference was observed in control animals, thus corroborating the notion that the stimulation of IPACL^CRH^ neurons is perceived as an aversive stimulus that entails arousal and avoidance behaviour (Fig. 1, J **and** K).

To comprehend the efferent projections of IPACL^CRH^ neurons, we injected a Cre-dependent AAV into extended amygdala nuclei of CRH-Cre::Ai9 reporter mice to express the presynaptic marker synaptophysin-GFP (Syn-GFP) in CRH^+^ neurons. Within the extended amygdala, IPACL^CRH^ neurons presented a distinct pattern of distant projections (fig. S3, A **to** F **and Movie S3**). While efferents of GABAergic IPACL CRH^-^ neurons were detectable throughout the SN and VTA (Fig. 2A), terminals of IPACL^CRH^ neurons were specifically confined to the SN possessing a high CRH content as detected by immunohistochemistry (Fig. 2, B **and** C **and** fig. S3, G **and** H **and Movie S4**). Next, we injected the retrograde tracer FluoroGold^TM^ (FG) into the SN of CRH-Cre::Ai9 reporter mice. Retrogradely labelled cells were detected in all brain regions previously shown to innervate the SN (*23*). Additionally, and in line with anterograde viral tracing, we observed a substantial number of FG labelled CRH neurons in the CeA and IPACL (Fig. 2, D **to** F). These results raised the intriguing question of whether the prominent IPACL-SN projections are sufficient to convey the behavioural response following stimulation of IPACL^CRH^ neurons.

**Figure 2:**
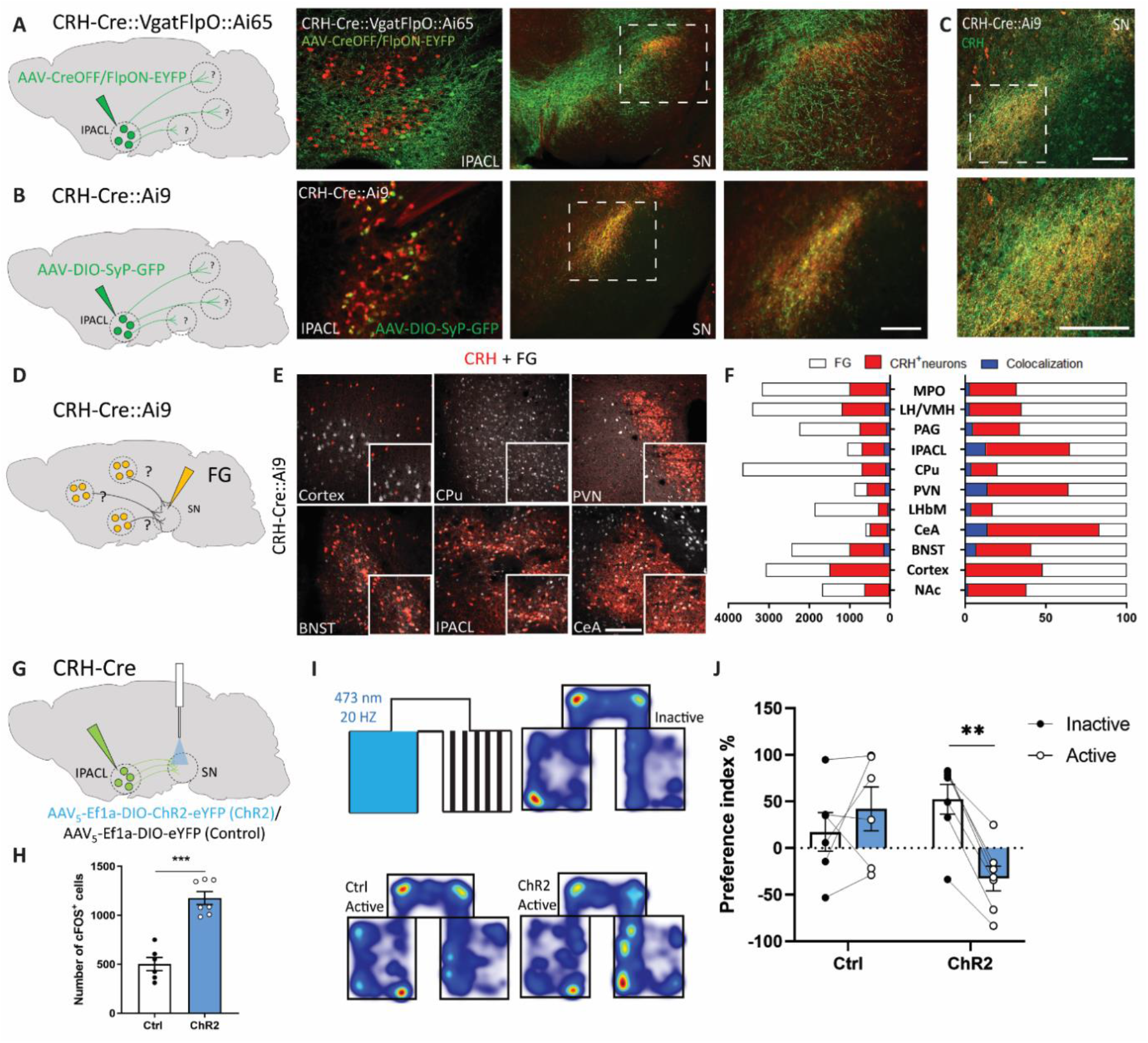
Projections of IPACL^CRH^ neurons to the SN promote avoidance behaviour. **A**, Scheme illustrating injection of AAV_DJ_-hSyn-CreOff/FlpOn-EYFP into IPACL of CRH-Cre::Vgat-FlpO::Ai65 mice. Representative image of the injection sites and respective target area in the SN. **B**, Scheme illustrating injection of AAV_9_-CMV-DIO-Syp-GFP into IPACL of CRH-Cre::Ai9 mice. Representative images of the injection site and respective target area in the SN. **C**, Representative images of CRH staining in the SN of CRH-Cre::Ai9 mice showing CRH expression in axon terminals. **D,** Scheme illustrating injection of FG into the SN of CRH-Cre::Ai9 animals. **E**, Representative images of various brain regions demonstrating localization of FG in tdTomato-expressing CRH neurons. **F**, Quantification of FG labelled CRH neurons throughout the brain following injection into the SN of CRH-Cre::Ai9 mice. **G**, Scheme illustrating injection of AAV_5_-Ef1a-DIO-ChR2(H134R)-EYFP (ChR2) or AAV_5_-Ef1a-DIO-EYFP (Ctrl) into the IPACL and placement of optic fibre in the SN of CRH-Cre mice. **H**, Quantification of cFOS positive cells in the SN (*p* < 0.0001, t = 7.13). **I**, RTPP paradigm with heatmaps visualizing the cumulative presence of mice in the test compartments before and during light activation. **J**, Preference index (two-way ANOVA, Bonferroni’s multiple comparison test, *p* = 0.008, t = 3.7). Values represent mean ± SEM, * *p* < 0.05, ** *p* < 0.01, *** *p* < 0.0001. All scale bars = 200 µm. Abbreviations: LH/VMH, lateral/ventromedial hypothalamus; LHbM, lateral habenula, medial; MPO, median preoptic area; NAc, nucleus accumbens; PAG, periaqueductal grey; PVN, paraventricular nucleus of the hypothalamus.

To address this, we first established whether optogenetic activation of IPACL^CRH^ neurons can mimic the effects of their chemogenetic activation. Therefore, we subjected mice selectively expressing ChR2 in IPACL^CRH^ neurons (fig. S4, A **to** D) to a real-time place preference test (RTPP). Illumination of ChR2 expressing neurons directly in the IPACL resulted in significant avoidance of the light-paired chamber recapitulating the consequences of CNO-induced activation of IPACL^CRH^ neurons during the conditioning phase (fig. S4, E **and** F). Next, we performed the RTPP but exclusively illuminated terminals of IPACL^CRH^ neurons in the SN (Fig. 2, G **and** H). Similar to the IPACL stimulation, ChR2-expressing mice also responded with strong place aversion compared to control mice (Fig. 2, I **and** J). These findings clearly indicate that the IPACL-SN projection is sufficient to drive the observed behavioural activation and avoidance. The direct impact of IPACL^CRH^ neurons on the SN is further supported by the observation that the CNO-induced activation of IPACL^CRH^ neurons entailed enhanced cFOS expression in the SN (fig. S4G).

To understand whether the behavioural consequences of IPACL^CRH^ neuron activation can be directly attributed to CRH release and downstream activation of CRH receptors, we repeated the behavioural paradigms while blocking the high affinity CRHR1 using the specific small molecule antagonist R121919. Systemic treatment of hM3Dq-expressing mice with R121919 was sufficient to attenuate the CNO-induced increase in cFOS expression in the SN (fig. S4H) and blocked the CNO-induced increase in locomotor activity and entries to the centre zone in the OF (Fig. 3A). In the DaLi, CRHR1 antagonism reduced the CNO-induced increase of transitions into the lit zone without affecting the time spent in that compartment compared to mice treated with CNO only (Fig. 3B). In the CPP, R121919 treatment prevented the place avoidance caused by the CNO-induced activation of IPACL^CRH^ neurons during conditioning (Fig. 3, C **and** D). Taken together, our data indicate that activation of IPACL^CRH^ neurons triggers arousal and avoidance behaviour in a CRHR1-dependent manner. Next, we investigated the SN innervation by IPACL^CRH^ neurons in more detail by injecting a Cre-dependent AAV expressing Syp-mCherry into CRH-Cre::CRHR1^ΔEGFP^ mice allowing simultaneous visualization of CRH terminals and CRHR1 expressing neurons (*5*) (Fig. 3E). Terminals of IPACL^CRH^ neurons were mainly found in the SN pars compacta (SNpc) additionally extending into the dorsal aspects of the SN pars reticulata (SNpr). snRNA-Seq of nuclei isolated from the ventral midbrain of CRHR1-Cre::INTACT mice revealed the presence of CRHR1 in DA, GABAergic and glutamatergic neurons. Depending on their localization, SN^CRHR1^ neurons are predominantly dopaminergic (SNpc) or GABAergic (SNpr) (fig. S5, A **to** K). Accordingly, CRH terminals were found interspersed between dopaminergic and GABAergic CRHR1^+^ neurons in the SNpc and dorsal SNpr (fig. S6, A **and** B).

**Figure 3:**
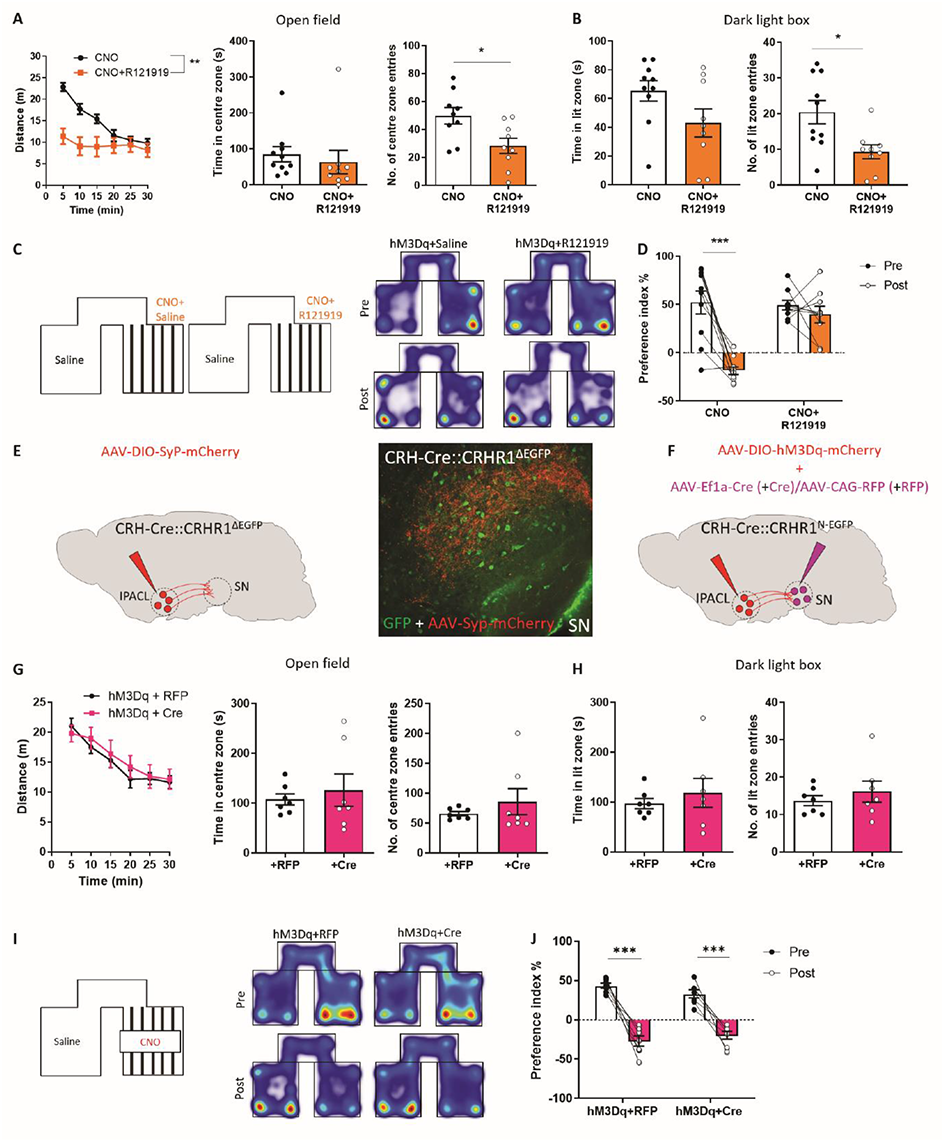
IPACL^CRH^ neurons modulate arousal and avoidance behaviour in a CRHR1-dependent manner. **A-D**, Treatment with the CRHR1 antagonist R121919 (10 mg/kg) blocks CNO-induced arousal and avoidance behaviour in mice expressing hM3Dq in IPACL^CRH^ neurons. **A**, Distance travelled (n = 9-10, repeated measures two-way ANOVA, *F*_1,17_ = 8.97, *p* = 0.0081), time in centre (*p* = 0.57, t = 0.56) and entries into centre zone (*p* = 0.017, t = 2.66) of the open field. **B**, Time spent in lit zone (*p* = 0.12, t = 1.64) and entries into lit zone (*p* = 0.022, t = 2.54) of the dark-light box. **C**, Schematic illustration of place preference paradigm and heat maps showing the accumulated time animals stayed in respective chambers pre and post conditioning. **D**, Calculated preference index (two-way ANOVA, Bonferroni’s multiple comparison, *p* < 0.0001, t = 5.94). **E**, Scheme illustrating injection of AAV_8_-CMV-DIO-Syp-mCherry into the IPACL of CRH-Cre::CRHR1^ΔEGFP^ mice. Representative images showing terminals of IPACL^CRH^ neurons in the SN. **F**, Scheme illustrating co-injection of AAV_8_-hSyn-DIO-hM3Dq-mCherry into the IPACL and AAV_1_-Ef1a-Cre or AAV_1_-CAG-RFP into the SN of CRH-Cre::CRHR1^N-EGFP^ mice. **G-J**, CNO treatment does not affect arousal or avoidance behaviour in hM3Dq-expressing animals. **G**, Distance travelled (repeated measures two-way ANOVA, *F*_1,12_ = 0.12, *p* = 0.74), time in centre zone (*p* = 0.59, t = 0.55) and entries into centre zone (*p* = 0.38, t = 0.91) of the OF. **H**, Time in lit zone (*p* = 0.49, t = 0.69) and entries into lit zone (*p* = 0.45, t = 0.77) of the DaLi. **I**, CPP paradigm with heatmaps visualizing the cumulative presence of mice in the test compartments pre and post CNO treatment. **J**, Preference index (repeated measures two-way ANOVA, *F*_1,12_ = 108.2, *p* < 0.0001). Values represent mean ± SEM, * *p* < 0.05, ** *p* < 0.01, *** *p* < 0.0001. All scale bars = 200µm.

To directly investigate the role of SN^CRHR1^, we activated IPACL^CRH^ neurons and simultaneously ablated CRHR1 from neurons in the SN. To this end, we expressed hM3Dq in IPACL^CRH^ neurons and injected AAV-Ef1a-Cre (+Cre) or AAV-CAG-RFP (+RFP) into the SN of CRH-Cre::CRHR1^N-EGFP^ mice possessing a GFP tagged CRHR1, which is sensitive to deletion by Cre recombinase (*5*) (Fig. 3F **and** fig. S6C). However, Cre-injected animals lacking CRHR1 in SN neurons were indistinguishable from control mice in all tested paradigms (Fig. 3, G **to** J). These results suggest that IPACL^CRH^ neurons convey arousal and avoidance behaviour largely independent of CRHR1 expression in SN neurons. Along these lines, chemogenetic activation of SN^CRHR1^ neurons did not reproduce the behavioural activation and arousal triggered by stimulation of IPACL^CRH^ neurons (fig. S7, A-G).

These findings beg the question to what extent IPACL^CRH^ neurons are synaptically connected with SN^CRHR1^ neurons and where CRH, potentially released following IPACL stimulation, is acting upon CRHR1? Therefore, we performed rabies virus-assisted retrograde monosynaptic tracing injecting pseudotyped rabies virus into the SN of CRHR1-Cre animals specifically expressing TVA in SN^CRHR1^ neurons (fig. S8A). Transsynaptic tracing revealed that SN^CRHR1^ neurons are innervated by numerous brain structures but eminently by the CeA, BNST, caudate putamen (CPu) and external globus pallidus (GPe), whereas the IPACL showed only minor direct synaptic contacts with SN^CRHR1^ neurons (fig. S8, B, C **and** F). To gain a more comprehensive view of SN afferent connections, we performed rabies virus tracing injecting Flp-dependent AAV-CBh-fDIO-TVA-2A-GFP-OG into Vgat-FlpO (SN^GABA^ neurons) or TH-FlpO mice (SN^DA^ neurons) (fig. S8A). The majority of neurons projecting to SN^DA^ and SN^GABA^ neurons were detected in the CPu and the GPe, structures which are largely devoid of CRH expressing neurons (fig. S8, B **and** D, E **and** G, H). However, the CPu and particularly the GPe are rich in CRHR1-expressing neurons (*24*). As a major component of the basal ganglia circuit, the GPe is strongly interconnected with the SN. We observed that a large proportion of GPe^CRHR1^ neurons co-express PV (fig. S9, A **and** B) and anterograde tracing by injecting AAV-Ef1a-DIO-Syp-mCherry into the GPe of CRHR1-Cre animals demonstrated that GPe^CRHR1^ neurons send strong direct efferents into the SNpr (**Movie S5**). Specifically, these GABAergic GPe^CRHR1/PV^ long-range projection neurons show strong innervation of SNpr^PV^ neurons in the SN (fig. S9, C **and** D). This was further substantiated using retrograde tracing by injecting pseudotyped rabies virus into the SN of PV-Cre animals resulting in predominant labelling of neurons in the GPe and CPu (fig. S9, E **and** F).

These findings raise the intriguing possibility that presynaptic CRHR1 located on GPe^CRHR1^ efferents is the principal recipient of CRH released from IPACL^CRH^ neuron terminals in the SN. Therefore, we probed whether stimulation of GPe^CRHR1^ neurons is capable of affecting arousal and avoidance behaviour. We observed that locally confined optogenetic stimulation of GPe^CRHR1^ terminals indeed triggered place avoidance in mice expressing ChR2 in GPe^CRHR1^ neurons (fig. S10, A **to** D). Similarly, chemogenetic activation of GPe^CRHR1^ neurons in CRHR1-Cre animals triggered arousal and avoidance behaviour (fig. S11, A **to** G). To further corroborate an interaction between these circuits, we investigated the convergence of presynaptic terminals originating from IPACL^CRH^ and GPe^CRHR1^ neurons within the SN. Application of AAV-hSyn-CreOff/FlpOn-EYFP into the IPACL and AAV-hSyn-DIO-mCherry into the GPe of CRH-FlpO::CRHR1-Cre animals revealed that axon terminals of IPACL^CRH^ and GPe^CRHR1^ neurons are located in close proximity within the SN enabling activation of presynaptic CRHR1 by CRH released from IPACL^CRH^ neuron terminals (Fig. 4A).

**Figure 4:**
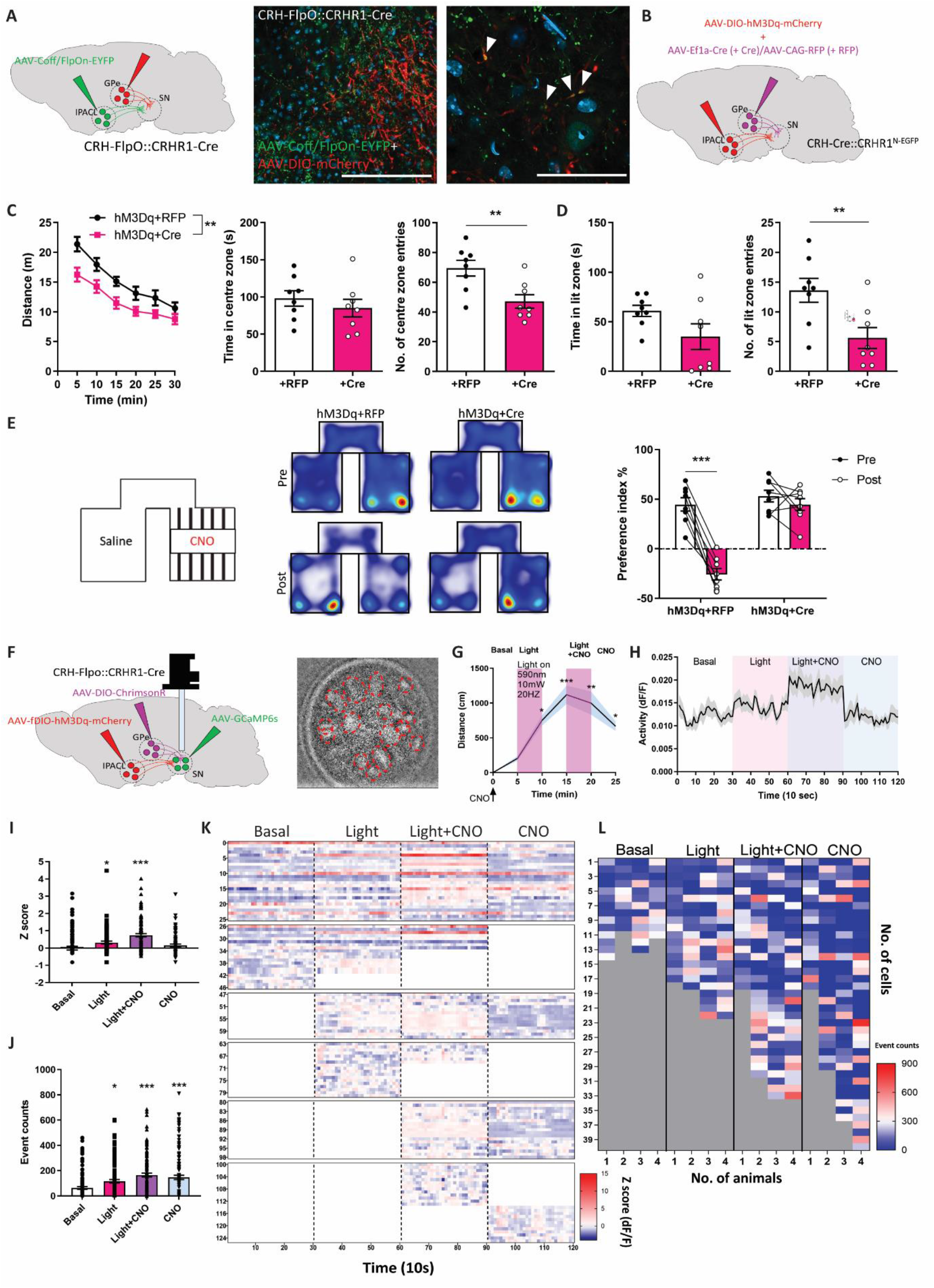
CRHR1 on GPe^CRHR1^ neuron terminals is required for arousal and avoidance behaviour driven by IPACL^CRH^ neurons. **A**, Scheme displaying injection of AAV_DJ_-hSyn-CreOff/FlpOn-EYFP into the IPACL and AAV_8_-hSyn-DIO-mCherry into the GPe of CRH-FlpO::CRHR1-Cre mice. Representative images showing terminals of IPACL^CRH^ and GPe^CRHR1^ neurons in the lateral SN, scale bar = 100 µm (right panel). White arrowheads indicate co-localization of IPACL^CRH^ and GPe^CRHR1^ neuron terminals. **B**, Scheme illustrating co-injection of AAV_8_-hSyn-DIO-hM3Dq-mCherry in the IPACL and AAV_1_-Ef1a-Cre or AAV_1_-CAG-RFP in the GPe of CRH-Cre::CRHR1^N-EGFP^ mice. **C-E**, Ablation of CRHR1 from GPe^CRHR1^ neurons reduces arousal and avoidance behaviour induced by CNO treatment in hM3Dq-expressing animals. **C**, Distance travelled (repeated measures two-way ANOVA, *F*_1,14_ = 9.26, *p* = 0.0088), time in centre zone (*p* = 0.42, t = 0.83) and entries into centre zone (*p* = 0.006, t = 3.21) of the OF. **D**, Time in lit zone (*p* = 0.087, t = 1.84) and entries into lit zone (*p* = 0.0095, t = 3) of the DaLi. **E**, CPP paradigm with heatmaps visualizing the cumulative presence of mice in the test compartments pre and post CNO treatment. Preference index (two-way ANOVA, Bonferroni’s multiple comparison test, *p* < 0.0001, t = 8.23). **F**, Scheme displaying injection of AAV_8_-hSyn-fDIO-hM3Dq-mCherry into the IPACL, AAV_8_-hSyn-DIO-ChrimsonR-tdTomato into the GPe and AAV5-hSyn-GCaMP6s into the SN as well as placement of GRIN lens above the lateral SN of CRH-FlpO::CRHR1-Cre mice. Representative image showing cells with fluorescence signal as detected by the miniscope (encircled by red dashed lines). **G**, Locomotor activity of mice subjected to different stimulation regimes (one-way ANOVA, Bonferroni’s multiple comparison test, Light, *p* = 0.01; CNO, *p* < 0.0001; Light+CNO, *p* = 0.002; CNO, *p* = 0.036). **H**, Representative activity traces of a cluster of cells. **I**, Cell activity under different stimulation regimes compared to basal conditions (one-way ANOVA, Bonferroni’s multiple comparison test, Light, *p* = 0.033; Light+CNO, *p* < 0.0001; CNO, *p* = 0.265). **J**, Event counts following different stimulations compared to basal conditions (one-way ANOVA, Bonferroni’s multiple comparison test, Light, *p* = 0.027; Light+CNO, *p* < 0.0001; CNO, *p* = 0.0001). **K**, Heatmap visualization of cell activities active during different conditions. **L**, Heatmap visualization of event counts during different conditions. Values represent mean ± SEM, * *p* < 0.05, ** *p* < 0.01, *** *p* < 0.0001. All scale bars = 200 µm.

To test this hypothesis, we activated IPACL^CRH^ neurons and simultaneously deleted CRHR1 from GPe^CRHR1^ neurons using CRH-Cre::CRHR1^N-EGFP^ mice injected with AAV-hSyn-DIO-hM3Dq in the IPACL and AAV-Ef1a-Cre in the GPe (Fig. 4B **and** fig. S12, A **to** C). The ablation of CRHR1 from GPe^CRHR1^ neurons was able to prevent the behavioural arousal and place avoidance behaviour previously seen following activation of IPACL^CRH^ neurons (Fig. 4, C **to** E **and** fig. S12, D **and** E). In contrast, neither deletion of CRHR1 from GPe^CRHR1^ neurons per se nor unspecific effects of Cre expression entailed any behavioural alterations (fig. S12, F **to** H). To rule out that GPe^CRHR1^ neurons receive direct synaptic input from IPACL^CRH^ neurons, we conducted rabies tracing in the GPe of CRHR1-Cre animals. While retrogradely labelled cells were observed in multiple brain regions, we only occasionally detected isolated mCherry^+^ cells in the IPACL (fig. S13, A **to** C).

To interrogate the connectivity of the IPACL-SN-GPe circuit functionally, we performed in vivo calcium imaging in the SN while stimulating either IPACL^CRH^ neurons or GPe^CRHR1^ terminals. We expressed ChrimsonR in the GPe, hM3Dq in the IPACL and GCaMP6s in the lateral SN of CRH-FlpO::CRHR1-Cre animals, which were subjected to a 25 min OF with continuous recording (Fig. 4F **and** fig. S14A). Mice showed behavioural activation following combined CNO and light application as evidenced by increased locomotor activity (Fig. 4G). We recorded 125 cells in total, with specific populations of cells (50%) remaining active during different conditions and cells, which were recruited during different phases of stimulation (Fig. 4H **and** fig. S14B **to** E). We observed significantly increased neuronal activity by activation of GPe^CRHR1^ terminals (Light) as well as by activation of both IPACL^CRH^ neurons and GPe^CRHR1^ terminals (Light+CNO) (Fig. 4, I **and** K). Activation of IPACL^CRH^ neurons alone (CNO) did not induce any differences compared to basal conditions potentially due to exhaustion of the sensor at the end of the imaging session. However, we detected significantly more event counts for all three experimental conditions compared to baseline (Fig. 4, J **and** L). Taken together, our calcium imaging data support a functional link between IPACL^CRH^ and GPe^CRHR1^ neurons governing neuronal activity in the SN.

Here we identified a tripartite CRH circuit consisting of GABAergic IPACL^CRH^ and GPe^CRHR1^ neurons whose efferents converge in the SN. Stimulation of IPACL^CRH^ neurons is perceived as aversive, entailing arousal and avoidance behaviour, which adds to the recently described role of the IPAC in controlling aversive responses in the context of innate and learned disgust (*25*). Presynaptically localized CRH receptors and the capacity of presynaptic CRHR1 to facilitate GABA release has been demonstrated (*26–28*). Accordingly, our data suggest that GABA release from GPe^CRHR1^ neurons is triggered by direct stimulation and further facilitated by presynaptic CRHR1 activated by CRH released from IPACL^CRH^ efferents in the SN. We found that the majority of GPe^CRHR1^ neurons express PV. This subpopulation of GPe neurons has been shown to directly innervate the SNpc and SNpr (*29–31*). Along these lines, we observed that GPe^CRHR1^ neurons predominantly form synapses at perikarya and primary dendrites of inhibitory SNpr^PV^ cells, which represent a major output of the basal ganglia circuit (*32*). Optogenetic or chemogenetic activation of SNpr basal ganglia output neurons has been shown to impair spontaneous locomotion and to block active avoidance (*33, 34*). In contrast, inhibition of SNpr neurons facilitates avoidance behaviour (*34*). This is in agreement with our observation that stimulation of IPACL^CRH^ or GPe^CRHR1^ neurons, likely resulting in inhibition of SNpr projection neurons, promotes avoidance behaviour in the applied place preference paradigms. In parallel, enhanced inhibition of GABAergic output neurons might also disinhibit the inhibitory drive on SNpc^DA^ neurons, which is conveyed via axon collaterals of SNpr output neurons (*35*). It remains to be investigated to what extent stimulation of IPACL^CRH^ neurons directly affects DA release, which might further contribute to the observed hyperlocomotion. Only recently, another CRH-specific circuit has been described connecting the CeA and the lateral SN contributing to emotional behaviours involving appetitive and aversive learning (*36*). The SN has classically been regarded as part of the basal ganglia motor circuit controlling voluntary movements. However, accumulating evidence including our findings contradict a clear anatomical and functional segregation of midbrain DA systems (*37*). Finally, it is tempting to speculate that this novel extended amygdala – basal ganglia circuit might contribute to pathological hyperarousal and maladaptive avoidance responses, which are hallmarks of anxiety and depression.

## Acknowledgements

We would like to thank Albin Varga, head of the animal facility, and his staff for their dedicated support with animal care; Stefanie Unkmeir, Sabrina Bauer and the scientific core unit Genetically Engineered Mouse Models for genotyping support; Elfi Fesl and Jessica Keverne for proof reading and editing of the manuscript. We thank Carsten Wotjak for his input and critical discussion; Magdalena Götz, Silvia Cappello and Karl-Klaus Conzelmann for the generous technical support.

## Funding

Max Planck Society (JMD)

German Ministry of Science and Education, IMADAPT, FKZ 01KU1901 (JMD)

German Research Foundation, project-ID 118803580, SFB 870 Z1 (AHH)

Marie Skłodowska-Curie innovative training network PurinesDX (JMD)

## Author contributions

Conceptualization: SC, JMD Methodology: FF, CLL, AAH, RK

Investigation: SC, FF, CLL, LH, MJ, RDG, MG, DM

Data analysis: SC, LH, RDG, MZ, ME, NC, JMD

Visualization: SC, LH, RDG, ME, JMD

Funding acquisition: JMD

Project administration: JMD

Supervision: MZ, ME, NC, JMD

Writing – original draft: JMD, SC

Writing – review & editing: SC, MJ, ME, NC, JMD

## Competing interests

The authors declare that they have no competing interests.

## Data and materials availability

All data are available in the manuscript or the supplementary materials. Materials are available by request from the corresponding author.

## Supplementary Materials

### Materials and Methods

#### Mice

All animal experiments were conducted with the approval of and in accordance to the Guide of the Care and Use of Laboratory Animals of the Government of Upper Bavaria, Germany. Mice were group housed under standard lab conditions (22 ± 1°C, 55 ± 5% humidity) and maintained under a 12 h light-dark cycle with food and water ad libitum. All experiments were conducted with adult male mice (age: 2-5 months). The following mouse lines were used: CRH-Cre (Jackson Laboratory stock no. 012704) (*38*), CRH-FlpO (Jackson Laboratory stock no. 031559) (*39*), CRHR1-Cre (*9*), CRHR1^Δ-EGFP^ (*5*), CRHR1^N-EGFP^ (*5*), PV-Cre (*40*), TH-FlpO (*41*), Vgat-FlpO (Jackson Laboratory stock no. 029591) (*42*), INTACT (Jackson Laboratory stock no. 021039) (*43*), Ai9 (Jackson Laboratory stock no. 007909) (*44*), Ai65 (Jackson Laboratory stock no. 021875) (*45*). The following double and triple transgenic lines were generated in this study by cross-breeding of single transgenic lines: CRH-Cre::Ai9, CRH-Cre::Vgat-FlpO::Ai65, CRH-Cre::CRHR1^Δ-EGFP^, CRH-Cre::CRHR1^N-EGFP^, CRH-FlpO::CRHR1-Cre, CRHR1-Cre::INTACT. Primer sequences and protocols for genotyping are available upon request.

#### Electrophysiology

8-12 week old CRH-Cre::Ai9 mice were anesthetized with isoflurane and decapitated. The brain was rapidly removed from the cranial cavity and, using a vibratome (HM650V, Thermo Scientific), 350 µm-thick coronal slices containing the IPACL were cut in an ice-cold carbogen gas (95% O_2_/5% CO_2_)-saturated solution consisting of (in mM): 87 NaCl, 2.5 KCl, 25 NaHCO_3_, 1.25 NaH_2_PO_4_, 0.5 CaCl_2_, 7 MgCl_2_, 10 glucose, and 75 sucrose. Slices were incubated in carbogenated physiological saline for 30 min at 34°C and, afterwards, for at least 30 min at room temperature (23-25°C). The physiological saline contained (in mM): 125 NaCl, 2.5 KCl, 25 NaHCO_3_, 1.25 NaH_2_PO_4_, 2 CaCl_2_, 1 MgCl_2_, and 10 glucose. In the recording chamber, slices were superfused with carbogenated physiological saline (2-3 ml/min flow rate) and CRH^+^ as well as CRH^-^ IPACL neurons were identified using epifluorescence and infrared videomicroscopy. Somatic whole-cell current- and voltage-clamp recordings (−70 mV holding potential, >1 GΩ seal resistance, <20 MΩ series resistance, 3 kHz low-pass filter, 15 kHz sampling rate) were performed at room temperature using an EPC 10 amplifier (HEKA). For current-clamp measurements, patch pipettes (3-5 MΩ open tip resistance) were filled with a solution consisting of (in mM): 135 KMeSO_4_, 8 NaCl, 0.3 EGTA, 10 HEPES, 2 Mg-ATP, and 0.3 Na-GTP. In these experiments, NBQX (5 µM) and picrotoxin (100 µM) were added to the physiological saline. Current injections were used to depolarize or hyperpolarize the neuron under investigation. For voltage-clamp recordings of spontaneous AMPA receptor-mediated excitatory postsynaptic currents (AMPA-sEPSCs), the pipette solution consisted of (in mM): 125 CsCH_3_SO_3_, 8 NaCl, 10 HEPES, 0.5 EGTA, 4 Mg-ATP, 0.3 Na-GTP, and 20 Na_2_-Phosphocreatine. In these experiments, picrotoxin (100 µM) was added to the physiological saline. For recordings of spontaneous GABA_A_ receptor-mediated inhibitory postsynaptic currents (GABA_A_-sIPSCs), the pipette solution was composed of (in mM): 140 KCl, 5 NaCl, 10 HEPES, 0.1 EGTA, 2 Mg-ATP, 0.3 Na-GTP, and 20 Na_2_-Phosphocreatine. In these experiments, NBQX (5 µM) was added to the physiological saline. Offline analysis of electrophysiological data was performed using the Fitmaster software (HEKA) and Igor Pro program (WaveMetrics).

#### Immunohistochemistry

Mice were sacrificed with an overdose of isoflurane (Floren, Abbott) and transcardially perfused with 20 ml phosphate buffered saline (PBS) followed by 20 ml 4% paraformaldehyde (PFA). Dissected brains were post-fixed in 4% PFA overnight at 4°C and 50 µm-thick sections were prepared using a vibratome (Microm HM 650 V, Thermo-Fisher Scientific). Sections were rinsed in PBS and incubated overnight at 4°C with the primary antibody in PBS with 0.5% Triton-X100. On the next day, sections were washed, incubated with the secondary antibody coupled to suitable fluorescence (1:250 Alexa Fluor, Invitrogen, Thermo-Fisher Scientific) and afterwards washed with PBS. Sections were mounted using Fluoromount-G mounting medium (SouthernBiotech) and either left to air-dry or stored at −20°C for image acquisition. Primary antibodies: chicken-anti GFP antibody (1:500, Aves), rabbit-anti TH antibody (1:1000, Millipore), mouse-anti PV antibody (1:500, Swant), rabbit-anti RFP antibody (1:500, Rockland), rabbit-anti cFOS antibody (1:1000, Abcam), rabbit-anti WFS1 (1:200, St John’s Laboratory), mouse-anti Calbindin (1:500, Sigma), rabbit-anti SOM (1:500, Peninsula, Laboratories International), rabbit-anti Calretinin (1:1000, SYSY) and mouse-anti Pkcƌ (1:200, BD Bioscience). For quantification, all slides were randomized and coded before quantitative analysis. TH, PV and cFOS labelled cells were counted on every sixth section through the entire rostro-caudal extent of regions of interest (ROIs, 6 sections per animal). Images were captured with either a Zeiss Axioplan2 fluorescence microscope and Axio Vision 4.5 software or a Zeiss inverted laser scanning confocal microscope and Zen software. For confocal imaging, a z-stack of pictures of areas of interest was obtained with 0.4–1.2 μm step size and 800 × 800 to 1,024 × 1,024 pixel picture size. Images were analysed with ImageJ (http://rsweb.nih.gov/ij/).

#### CLARITY

Perfused mouse brains were fixed in 4% PFA overnight and then transferred into 1% hydrogel solution (1% acrylamide, 0.0125% bis-acrylamide, 4% PFA, 0.25% VA-044 initiator) for 48 h. The samples were degassed (nitrogen replacing oxygen, using 50 ml caps with tubing for 30 min) and polymerized (overnight at 37°C) in a 50 ml tube. Brains were removed from hydrogel and washed with 200 mM NaOH-Boric acid buffer (pH 8.5) containing 8% SDS for 12 h at 37°C (without shaking) to remove residual PFA and monomers. Brains were transferred to a flow-assisted clearing device using a temperature-control circulator. Next, 100 mM Tris-Boric buffer (pH 8.5) containing 8% SDS was used to accelerate the clearing (at 37-40°C). After clearing, the brains are washed in PBS with 0.2% Triton-X100 for at least 48 h at 37°C to remove residual SDS. Brains were incubated in sorbitol refractive index matching solution (sRIMS) for several days at room temperature and then subjected to imaging using a LaVision Light Sheet microscope (LaVision BioTec, Duisburg, Germany).

#### Isolation and single nucleus RNA sequencing

Mice were sacrificed at 2 months of age by cervical dislocation. Brains were collected in ice cold PBS and processed for tissue dissection. The target region (ventral midbrain) was isolated by subjecting brains to a brain matrix (Fine Science Tools). Brains were cut into 1 mm thick slices covering the entire ROI. The dorsal part including the cortex was removed and the remaining tissue was homogenized in 2 ml of lysis buffer (0.32 M sucrose, 10 mM Tris pH 8.0, 5 mM CaCl2, 3 mM Mg acetate, 1 mM DTT, 0.1 mM EDTA, 0.1% Triton X-100) by douncing 10-50 times in a 7 ml dounce homogenizer. Lysate was transferred to a 4 ml ultracentrifugation tube (Beckman; 14 × 95 mm; 344061), and 2 ml of sucrose solution (1.8 M sucrose, 10 mM Tris pH 8.0, 3 mM Mg acetate, 1 mM DTT) was pipetted directly to the bottom of the tube. Ultracentrifugation was carried out at 24,400 rpm for 2.5 h at 4°C (Beckman; L8-70 M; SW80 rotor). After centrifugation, the upper two layers were removed by aspiration. The nuclei pellet was resuspended in PBS with 0.1% fetal bovine serum (FBS). Then samples were processed to fluorescence-activated cell sorting (FACS) using a FACSMelody^TM^ (Becton Dickinson) with a nozzle diameter of 100 μm and FACSFlow^TM^ medium. Debris and aggregated nuclei were gated out by forward and sideward scatter. Single nuclei were gated out by FSC-W/FSC-A. Gating for fluorophores was done using isolated nuclei possessing GFP in the nuclear membrane.

snRNA-Seq was performed by using 10× genomics kit following the instructions of the manufacturer. In brief, nuclei were mixed with solutions containing barcoded beads and partitioning oil. To achieve single nucleus resolution, nuclei were delivered at a limiting dilution. Next, gel beads were dissolved, primers were released, and any co-partitioned nucleus was lysed. Primers were mixed with the cell lysate and with a master mix containing reverse transcription reagents. This produced barcoded, full-length cDNA from polyadenylated mRNA. After incubation, products were broken up and pooled fractions recovered. Silane magnetic beads were used to purify the first-strand cDNA from the post RT reaction mixture. Barcoded, full-length cDNA was amplified via PCR to generate sufficient mass for library construction. Enzymatic fragmentation and size selection were used to optimize the cDNA amplicon size. A Chromium Single Cell 3’ Gene Expression library comprises standard Illumina paired-end constructs beginning and ending with P5 and P7. The 16 bp 10× barcode and 12 bp unique molecular identifier (UMI) were encoded in Read 1, while Read 2 was used to sequence the cDNA fragments. Sample index sequences were incorporated as the i7 index read. TruSeq Read 1 and TruSeq Read 2 are standard Illumina sequencing primer sites used in paired-end sequencing.

#### Analysis of single nucleus RNA sequencing data

Single nuclei data was analysed using the python package SCANPY(*46*). Quality controls (QC), including nucleus QC and gene QC, were carried out on the raw dataset. The following measures were assessed: number of UMIs, number of expressed genes and the fraction of mitochondrial genes. Of the 492,928 nuclei and 31,053 genes in the raw dataset, 54 nuclei with gene counts greater than 3,500 and 476,198 nuclei that had gene counts fewer than 200 were removed. Of these, 11 nuclei were filtered out for having a fraction of mitochondrial genes higher than 0.04. Moreover, 17,614 genes that were detected in less than 20 nuclei were filtered out. After filtering, 16,665 nuclei and 13,439 genes remained for further analysis. To normalize feature and gene counts, the R package scran(*47*) was used. 4,000 highly variable genes were selected after data normalization, which were used for dimensionality reduction with principal component analysis (PCA) and clustering through embedding into a K Nearest Neighbours (KNN) graph using the louvain algorithm implemented in SCANPY. The clusters were visualised with the UMAP algorithm. To minimize the number of doublets and to ensure that nuclei did not cluster by the cell cycle phases, doublets were identified using the python package scrublet(*48*) and nuclei at different stages in cell cycle were recognized using marker genes(*49*). A Welch t-test was performed for differential gene expression between clusters. Cluster-specific marker genes were used to annotate each cluster.

#### Stereotactic surgeries

Mice were anesthetized for surgery with isoflurane (Floren, Abbott), 2% v/v in O_2_ and placed in a stereotaxic frame (Kopf Instruments). Body temperature was maintained with a heating pad. Metacam (5 mg/kg bodyweight) was administered as a systemic analgesic. Virus and tracer were delivered by using a 33-gauge microinjection needle with a 2.5 μl Hamilton syringe (Hamilton) coupled to an automated microinjection pump (World Precision Instruments Inc.) at 100 nl/min. Post-surgery recovery included Metacam supplementation (5 mg/kg, subcutaneous injection) for 3 d after surgery, with daily monitoring of food intake.

#### Viral injections and tracing analyses

For retrograde tracing, 0.5 µl of FluoroGold^TM^ (Fluorochrome, LLC) were injected unilaterally in the ROI: SN (AP: −3, ML: 1.65, DV: −4.13). After incubation for 7 days, mice were sacrificed for further analysis. For anterograde tracing, 0.5 ul of AAV_9_-CMV-DIO-Synaptophysin-GFP or AAV_8_-Ef1a-DIO-Synaptophysin-mCherry (∼10^13^ vg/ml, MIT Vector Core) or AAV_DJ_-hSyn-CreOff/FlpOn-EYFP (∼10^12^ vg/ml, UNC vector core, kindly provided by the Deisseroth lab) were injected into BSTLD (AP: 0.14, ML: 0.9, DV: −3.9), BSTLP (AP: 0.14, ML: 0.9, DV: −4.3), CeA (AP: −1.22, ML: 2.8, DV: −4.7), IPACL (AP: 0.26, ML: 2, DV: −5) or GPe (AP: −0.34, ML: 1.8, DV: −4). Viral tracers were incubated for a minimum of 2 weeks before the mice were sacrificed.

For retrograde rabies virus tracing, we first injected AAV_1_-CBh-DIO-TVA-T2A-GFP-OG or AAV_1_-hSyn-fDIO-revTVA-t2A-GFP-OG (∼10^16^ vg/ml) into SN or GPe. Two weeks later, SAD-EnVA-dG-mCherry rabies virus was injected into the same region. One week after injection of rabies virus, mice were sacrificed for further analysis.

For chemogenetic experiments, 0.5 ul of virus AAV_8_-hSyn-DIO-mCherry or AAV_8_-hSyn-DIO-hM3Dq-mCherry (Addgene plasmid #50459 and #44361, ∼10^8^ vg/mL) were injected bilaterally into target regions (IPACL, SN or GPe) using previous coordinates. For optogenetic manipulation, 0.5 μl of AAV_5_-Ef1a-DIO-ChR2-EYFP or AAV_5_-Ef1a-DIO-EYFP virus (∼10^12^ vg/ml, UNC Vector Core, kindly provided by the Deisseroth lab) were injected into the IPACL. For in vivo calcium imaging experiments AAV_8_-hSyn-fDIO-hM3Dq-mCherry (∼10^16^ vg/mL) was injected into the IPACL, AAV_8_-hSyn-FLEX-ChrimsonR-tdTomato (∼10^12^ vg/mL, UNC Vector core)^48^ in the GPe and AAV_2/1_-hSyn-GCaMP6s (∼10^12^ vg/mL, UPenn Vector core) in the SN. Generally, mice were allowed to recover for at least for two weeks before entering behavioural experiments.

#### Placement of optic fibre

Two weeks after virus injection, an optic fibre (200 μm core, 0.39 NA, 1.25 mm ferrule, Thorlabs) was implanted bilaterally above the IPACL (AP: 0.26, ML: 2, DV: −4.7 mm) or SN (AP: −3, ML: 1.65, DV: −3.8 mm). Implants were secured with cyanoacrylic glue (Braun), and the exposed skull was covered with dental acrylic (Paladur, Heraeus). The animals were left for two weeks for recovery before subjecting them to the real time place preference test.

#### Behavioural assays

Mice were single housed for a week and entered the respective test apparatus in random order. Mice rested for one day between each behavioural test, except in the conditioned place preference test. All experiments were analysed using the automated video-tracking system ANYmaze (Stoelting, Wood Dale, IL) and Ethovision (Noldus, Wageningen).

For chemogenetic activation of neurons, CNO dihydrochloride was dissolved in 0.9% saline at a concentration of 2.5 mg/ml. CNO was administered through i.p. injection (5 mg/kg) 15 min before mice entered the behavioural test apparatus. For blockade of CRHR1, R121919(*50*) hydrochloride was dissolved in 0.9% saline with 5% of ethanol at a concentration of 1.25 mg/ml. R121919 was administered through i.p. injection (10 mg/kg) 30 min prior to each behavioural test.

The OF test was conducted in an evenly illuminated (< 15 lx) square apparatus (40 cm (l) × 40 cm (w) × 60 cm (h)). The test duration was 30 or 90 min. The 15-min DaLi box test was performed in an apparatus consisting of a secure black compartment (10 cm (l) × 20 cm (w) x 35 cm (h), < 5 lx) and an aversive, brightly illuminated white compartment (10 cm (l) × 20 cm (w) x 35 cm (h), 400 lx) connected by a tunnel.

The CPP apparatus (T shape, 40 cm (l) × 40 cm (w) × 35 cm (h)) consisted of a starting box and two conditioning boxes made of black acrylic boards. The conditioning compartments (18 cm (l) × 20 cm (w) × 35 cm (h)), were separated by a smaller compartment (10 cm (l) × 20 cm (w) x 35 cm (h)) in the middle. The left compartment had mosaic walls and smooth flooring, the right compartment had black walls and thin-grid flooring. The experiment consisted of 3 phases: i) preconditioning, ii) conditioning and iii) post-conditioning phase. During the preconditioning phase (pre-test trial) (d1), each mouse was subjected to the middle compartment with free access to all compartments for 10 min. The conditioning phase (d2-d4) consisted of six 30 min training sessions that were carried out during morning and afternoon training. CNO (5 mg/kg, i.p.) was always paired with the less preferred compartment in the afternoon. On d2, mice were treated with saline (10 ml/kg, i.p.) in the morning training session and immediately paired with the preferred compartment with access blocked for 30 min. In the afternoon, mice were treated with CNO and immediately paired with the less preferred compartment for 30 min. The morning and afternoon training sessions were repeated for 3 d in total from d2-d4. In the post-conditioning phase (test trial, d5), each mouse was subjected to the middle compartment with free access to the other compartments in the absence of treatments for 10 min.

In the RTPP, mice were subjected to a place preference chamber (as described above) for 20 min. One counterbalanced side of the chamber was assigned as the stimulation side. The LED (Thorlabs, M470F3, 470 nm) was triggered on the basis of the location of the animal by using primmax software and controllers. In the photostimulated side of the arena, which was assigned based on the initial preference, mice received a 470-nm stimulation of 20 Hz with 10 mW of power at the fibre tips. Bilateral stimulation of freely moving animals was achieved using a fibreoptic rotary joint (FRJ_1 × 2i_FC-2FC, Doric). Behavioural data were recorded via a CCD camera.

#### GRIN lens implantation and baseplate fixation

The gradient index (GRIN) lens was implanted 1 week after viral injection (AAV_2/1_-hSyn-GCaMP6s) in the SN. Mice were anesthetized as described above. Debris was removed from the hole and a customized blunted 20G needle (1 mm in diameter) was slowly lowered down into the brain (AP: −3, ML: 1.65, DV: −3.95 mm). After retraction of the needle, a GRIN lens (ProView lens; diameter, 0.6 mm; length, ∼7.4 mm, Inscopix) mounted on a GRIN lens-holder was slowly implanted above the lateral SN. The exposed skull was covered with dental acrylic (Paladur, Heraeus). Two weeks after GRIN lens implantation, mice were anesthetized and placed in the stereotaxic frame. A baseplate (BPL-2, Inscopix) attached to a miniature microscope was positioned above the GRIN lens. The focal plane was adjusted until neuronal structures and GCaMP6s dynamics were observed. The baseplate was fixed using C&B-Metabond (Parkell).

#### In vivo calcium imaging

For simultaneous recording of GCaMP6s fluorescence signals and light activation of terminals in the SN, a miniature microscope system for integrated calcium imaging and optogenetics was used (nVoke 2.0, Inscopix). Mice were habituated to the miniscope 3 days before the behavioural experiment. The basal level of calcium activity was recorded in the first 5 min after CNO application. Then the effect of GPe^CRHR1^ neuron activity was measured by illuminating ChrimsonR-expressing terminals in the SN (590nm, 10mW, 20Hz) for 5 min. Another 5 min later, CNO reached its full effect on hM3Dq-expressing IPACL^CRH^ neurons. Thus, the recorded activity of cells in the next 5 min reflected both, light-induced activation of GPe^CRHR1^ and CNO-induced activation of IPACL^CRH^ neurons. Cell activities recorded in the last 5 min correspond to CNO-activated IPACL^CRH^ neurons. Image acquisition and behaviour were synchronized. For imaging data processing and analysis, we used the Inscopix data processing software (version 1.3.0).

#### Corticosterone measurement

Plasma corticosterone concentrations were measured as previously described(*5*) using a commercially available RIA kit (MP Biomedicals) according to the manufacturer’s instructions.

#### Statistical analysis

No statistical methods were used to pre-determine sample sizes. The numbers of samples in each group were based on those in previously published studies. The mean ± SEM was determined for each group. Figure generation and statistical calculation were conducted by Graphpad Prism 9 software. Two-tailed Student’s t test was performed for comparison between two groups with one variation. One-way ANOVA was used for one group with multiple variations. Two-way ANOVA was performed for two groups with multiple variations; Bonferroni’s post-hoc test was used for multiple comparisons. Repeated measures two-way ANOVA was performed with two groups with continuous variations (time), Bonferroni’s post hoc test was used for multiple comparisons. Differences were considered significant when p was smaller than 0.05.

---

**Figure S1:**
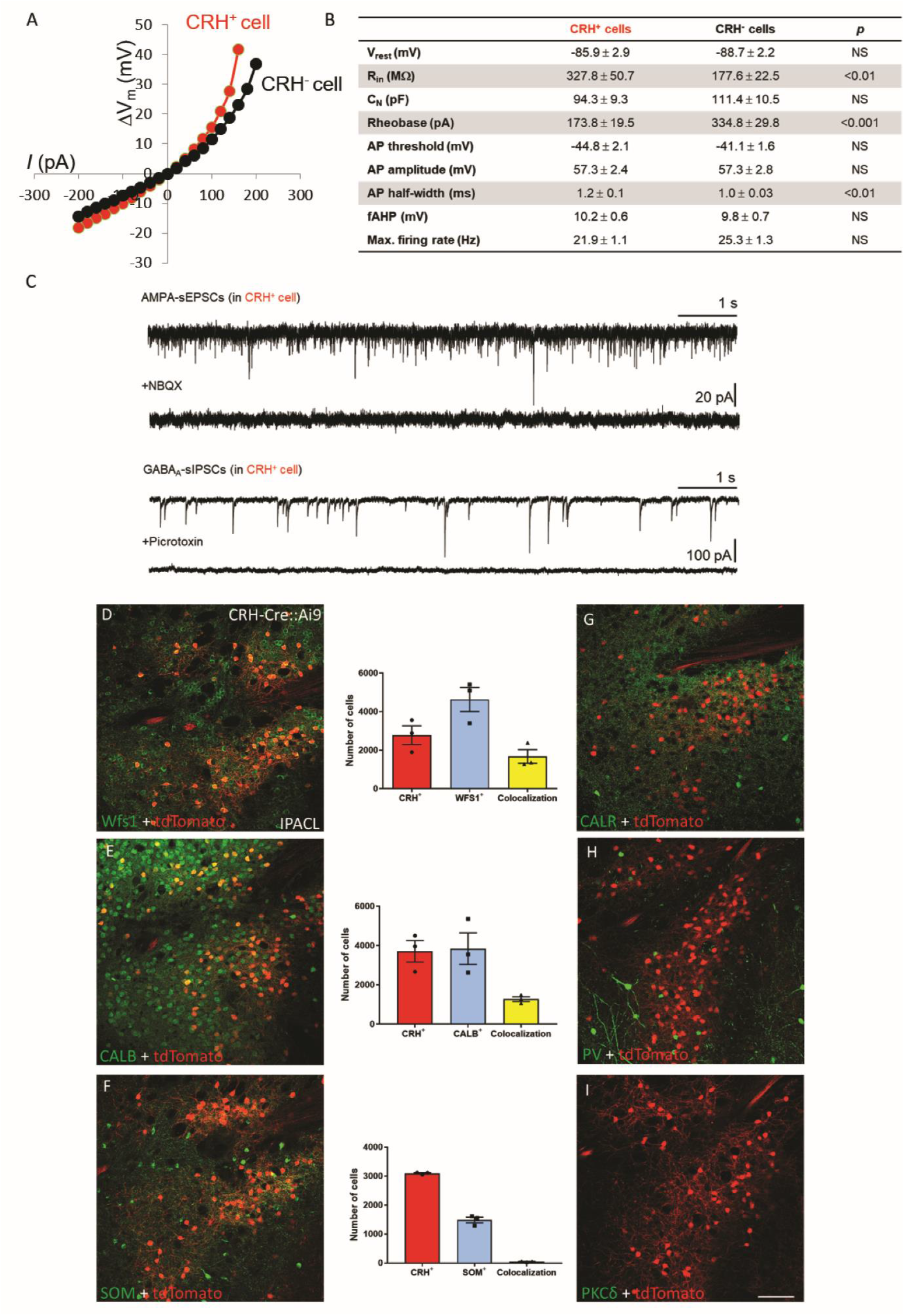
Characterization of CRH neurons in the extended amygdala. (**A**) Current-voltage relationship showing strong inward rectification in both CRH^+^ and CRH^-^ neurons. (**B**) Basic intrinsic properties of IPACL neurons. Both CRH^+^ (n = 15/5 mice) and CRH^-^ (n = 16/4 mice) cells have a remarkably low resting membrane potential (<-80 mV), which is typical for GABAergic spiny projection neurons. CRH^+^ neurons have higher input resistance and lower rheobase, indicating a higher excitability compared to CRH^-^ neurons. Abbreviations: V_rest_: resting membrane potential, R_in_: input resistance, C_N_: cell capacitance, AP: action potential, fAHP: fast after-hyperpolarization. Values represent mean ± SEM. Two-tailed unpaired t-test was used for statistical analysis. (**C**) IPACL^CRH^ neurons receive glutamatergic and GABAergic synaptic inputs. The upper representative recording traces show spontaneous AMPA receptor-mediated excitatory postsynaptic currents (AMPA-sEPSCs) in a CRH^+^ cell, which were completely blocked by NBQX. The lower traces demonstrate spontaneous GABA_A_-receptor-mediated inhibitory postsynaptic currents (GABA_A_-sIPSCs) in a CRH^+^ neuron, which totally disappeared in the presence of picrotoxin. (**D-I**) Representative images of IPACL^CRH^ neurons co-stained with markers of GABAergic neurons and quantification of co-localization. Abbreviations: CALB: calbindin, CALR: calretinin, PKCδ, protein kinase Cδ, PV: parvalbumin, SOM: somatostatin, WFS1: wolframin ER transmembrane glycoprotein 1. Values represent mean ± SEM, scale bar = 200 µm.

**Figure S2:**
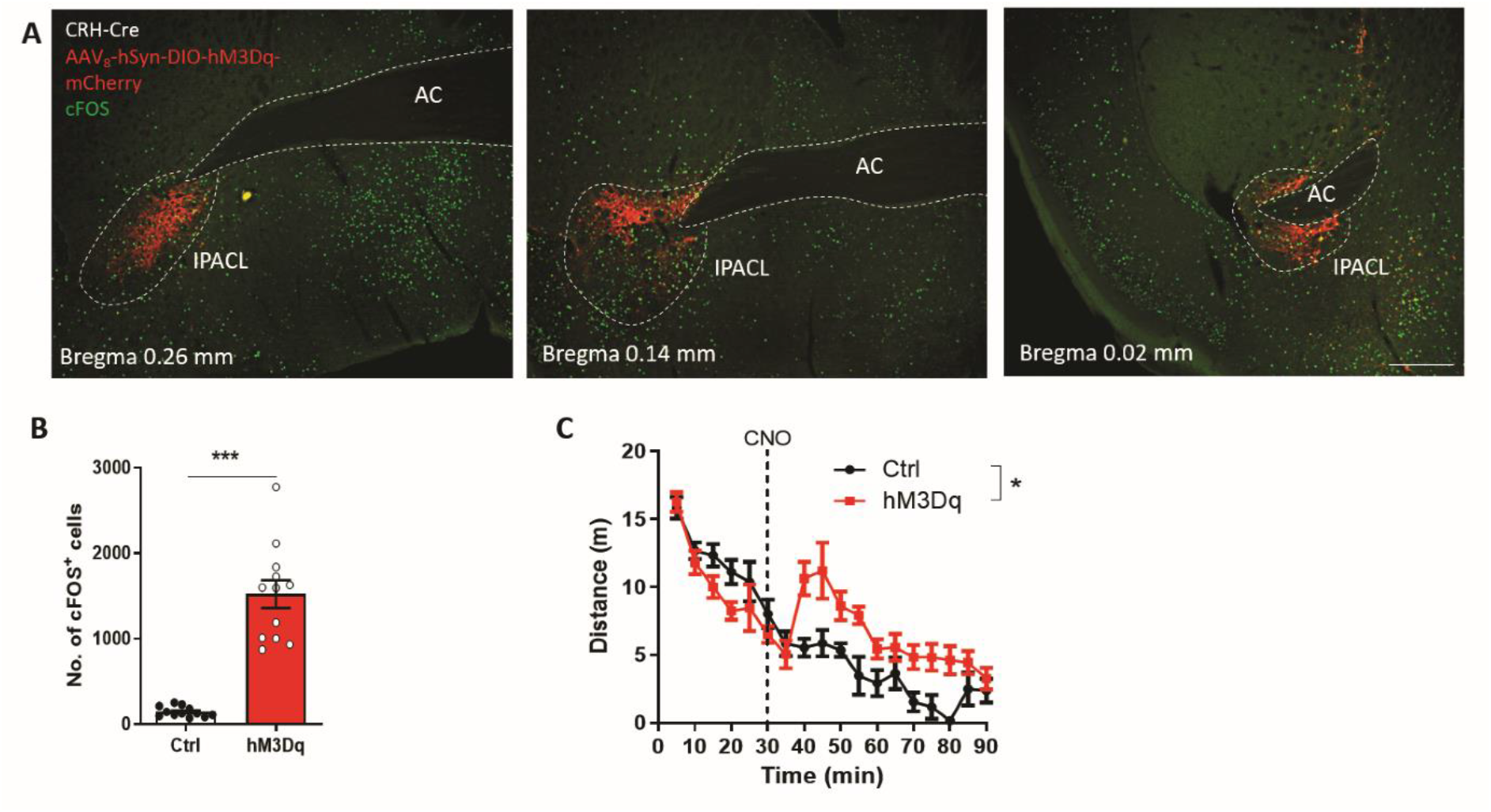
Chemogenetic activation of IPACL^CRH^ neurons. (**A)** Representative images of coronal brain sections of CRH-Cre animals injected with AAV_8_-hSyn-DIO-hM3Dq-mCherry (hM3Dq) in the IPACL and stained for cFOS following CNO administration. (**B**) Quantification of cFOS^+^ cells 90 min after CNO administration (*p* < 0.0001, t = 9.6). (**C**) Distance CRH-Cre mice expressing hM3Dq travelled in a 90-min open field test with CNO administration at 30 min (repeated measures two-way ANOVA, *F*_1,7_ = 6.28, *p* = 0.04).

**Figure S3:**
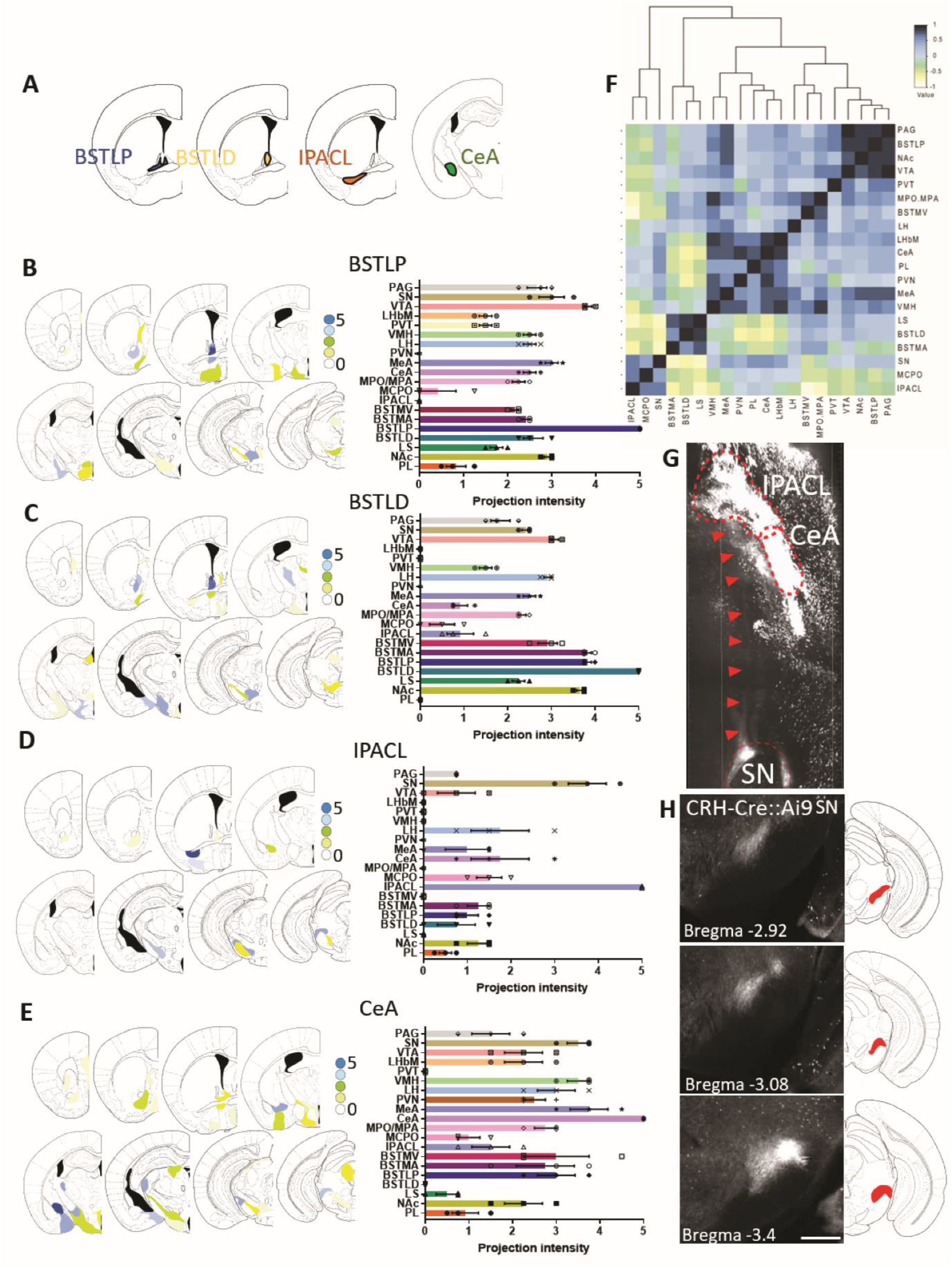
Anterograde tracing reveals target sites of CRH neurons of the extended amygdala. (**A**) Schematic illustration of injection sites of AAV_9_-CMV-DIO-Syp-GFP within extended amygdala subdivisions in CRH-Cre mice. (**B-E**) Summary of targets and strength of projections of CRH neurons located in the BSTLP, BSTLD, IPACL and CeA. (**F**) Pearson correlation demonstrating connectivity of brain regions based on anterograde tracing of CRH neurons. (**G**) Representative horizontal section of a cleared CRH-Cre::Ai9 brain showing projections (arrowheads) from the IPACL to the SN. (**H**) Coronal vibratome sections of a CRH-Cre::Ai9 brain reveals CRH neuron terminals in the SN. Values represent mean ± SEM, scale bar = 200 µm. Abbreviations: BSTLD, bed nucleus of the stria terminalis, laterodorsal; BSTLP, bed nucleus of the stria terminalis, lateroposterior; BSTMA, bed nucleus of the stria terminalis, medial, anterior; BSTMV, bed nucleus of the stria terminalis, medial, ventral; LH/VMH, lateral hypothalamus/ventromedial hypothalamus; LHbM, Lateral habenula, medial; LS, lateral septal nucleus; MCPO, magnocellular preoptic area; MeA, medial amygdaloid nucleus; MPO, median preoptic area; NAc, nucleus accumbens; PAG, periaqueductal grey; PL, prelimbic cortex; PVN, paraventricular nucleus of the hypothalamus; PVT, paraventricular thalamic nucleus.

**Figure S4:**
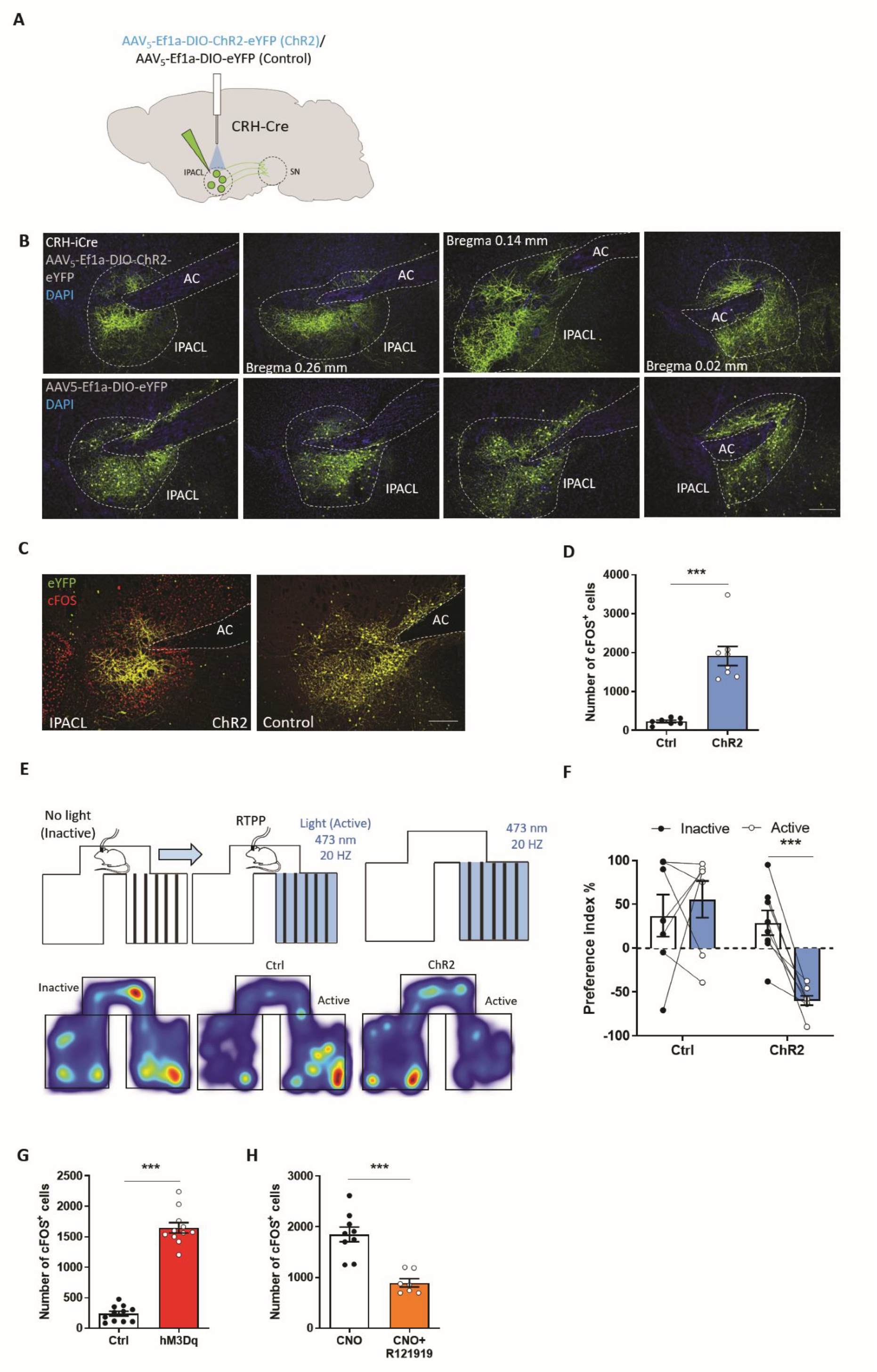
Optogenetic activation of IPACL^CRH^ neurons promotes arousal and avoidance behaviour. (**A**) Scheme illustrating injection of AAVs and placement of optic fibre in CRH-Cre mice. (**B**) Representative images of coronal brain sections of CRH-Cre animals injected with AAV_5_-Ef1a-DIO-ChR2-EYFP (ChR2) and AAV_5_-Ef1a-DIO-EYFP (Ctrl) in the IPACL. (**C**) Representative images of coronal IPACL sections stained for cFOS (90 min after light on). (**D**) Quantification of cFOS^+^ cells in the IPACL following light activation (*p* < 0.0001, t = 6.32). (**E**) Scheme depicting the real time place preference paradigm and heatmaps visualizing the cumulative presence of mice in the test compartments before and after light activation. (**F**) Preference index (two-way ANOVA, Bonferroni’s multiple comparison, *p* < 0.0001, t = 3.86). (**G**) Quantification of cFOS^+^ cells in the SN (*p* < 0.0001, t = 14.92) 90 min following CNO administration in mice expressing hM3Dq in IPACL^CRH^ neurons. (**H**) Quantification of cFOS^+^ cells in the SN (*p* = 0.0001, t = 5.3) 90 min following CNO + R121919 administration in mice expressing hM3Dq in IPACL^CRH^ neurons. Values represent mean ± SEM, * *p* < 0.05, ** *p* < 0.01, *** *p* < 0.0001, scale bar = 200 µm.

**Figure S5:**
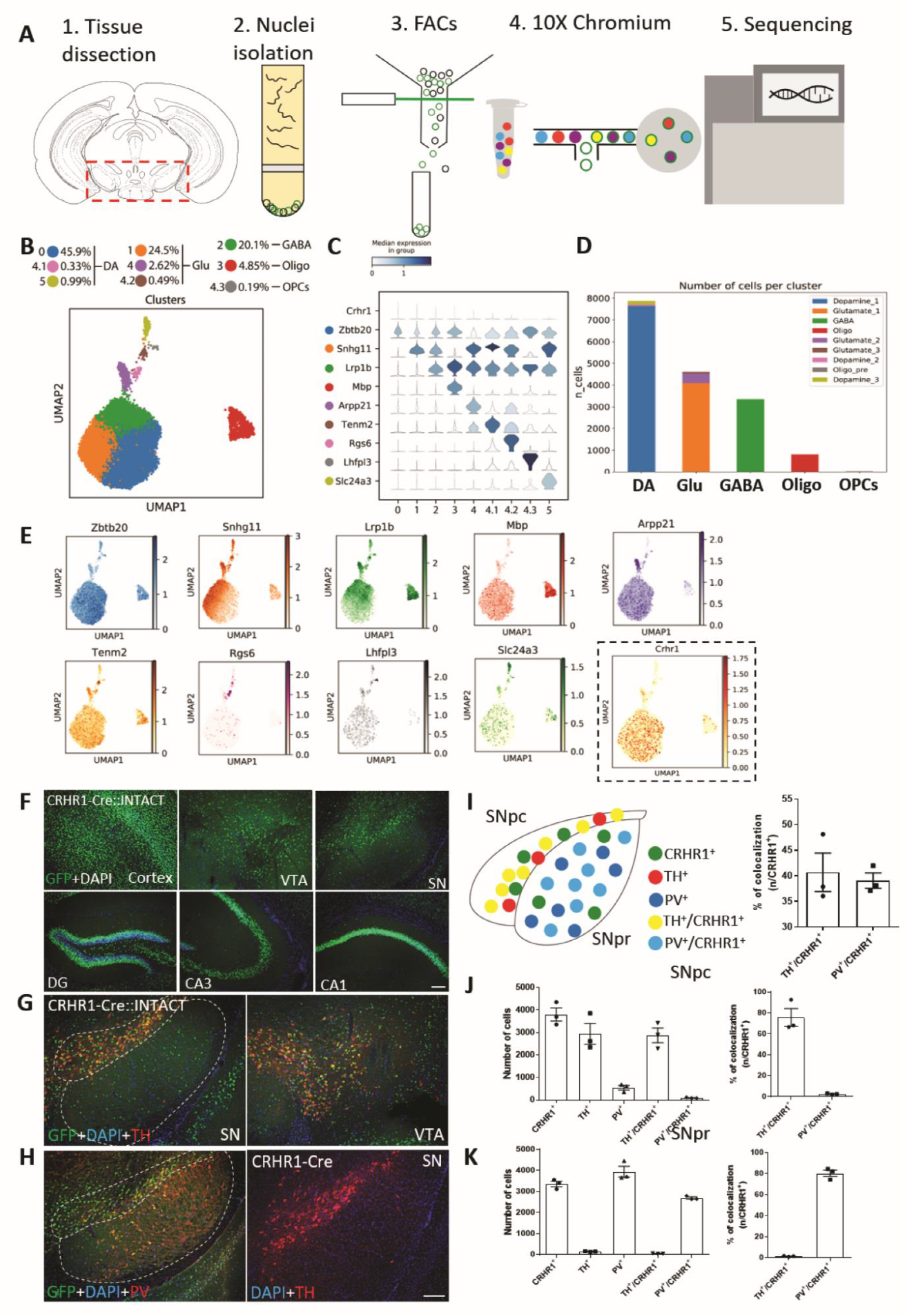
Single nucleus RNA sequencing (snRNA-Seq) using CRHR1-Cre::INTACT mice. (**A)** Scheme of workflow of snRNA-Seq in the ventral midbrain. (**B**) UMAP depicting 9 clusters of midbrain CRHR1 neurons, which can be specified into main clusters: dopaminergic (DA), glutamatergic (Glu) and GABAergic (GABA) CRHR1 neurons as well as oligodendrocytes (Oligo) and oligodendrocyte precursor cells (OPCs). (**C)** Violin plots depicting expression of marker genes for each cluster. (**D**) Distribution of marker genes for each cluster. (**E**) Number of cells belonging to different cell types. (**F**) GFP staining of nuclei of CRHR1 neurons in different brain regions of CRHR1-Cre::INTACT mice matching the expression pattern of endogenous CRHR1. (**G, H**) Co-staining of CRHR1 nuclei with markers of dopaminergic (TH) and GABAergic (PV) neurons in the SN. (**I**) Scheme illustrating the distribution of CRHR1 neurons within the SN and quantification of GABAergic and DAergic CRHR1 nuclei. (**J**) Distribution of DAergic and GABAergic CRHR1 neurons in the SN pars compacta (SNpc), and (**K**) in the SN pars reticulata (SNpr). Values represent mean ± SEM, scale bar = 200 µm.

**Figure S6:**
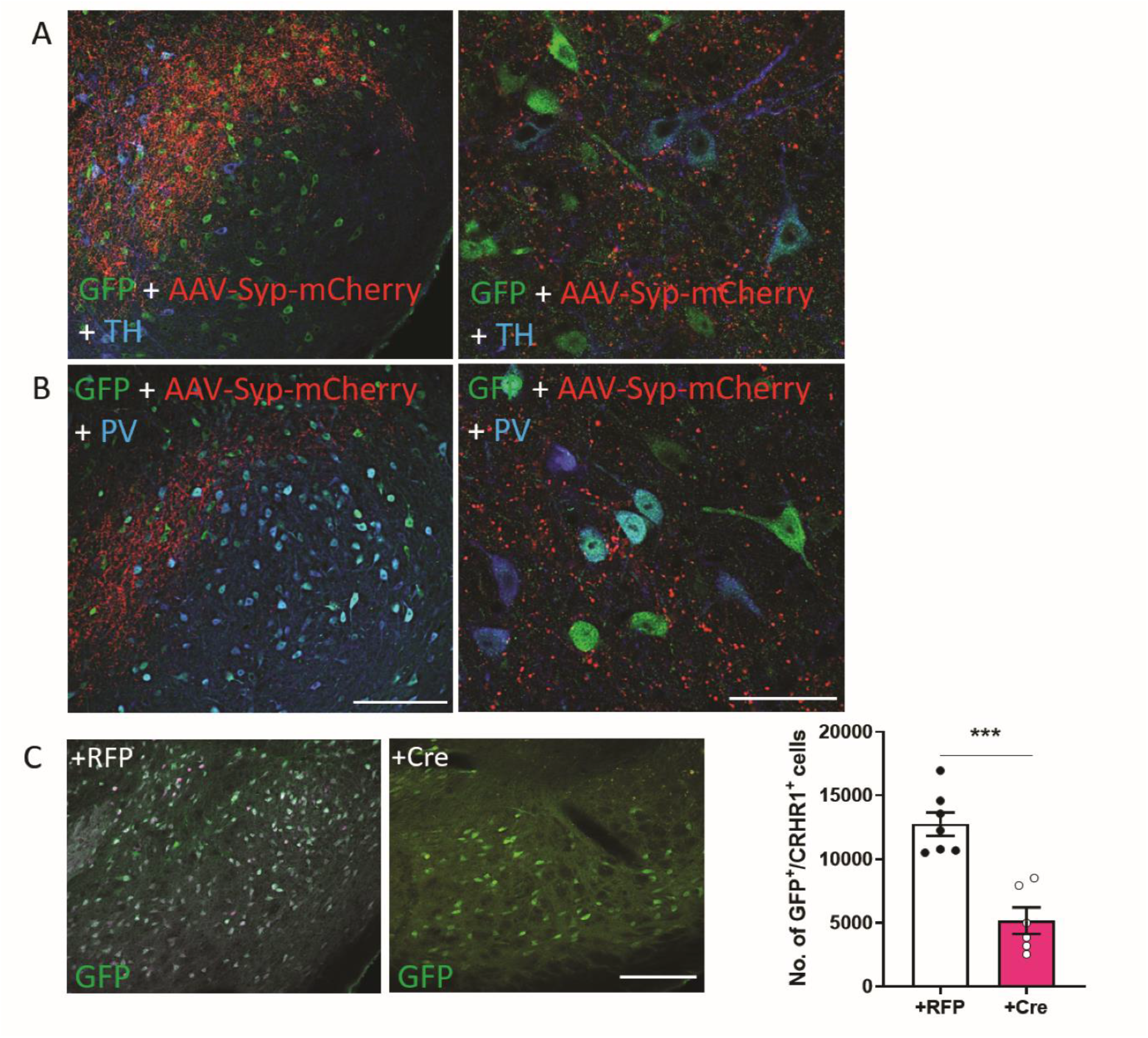
IPACLCRH axons are interspersed between CRHR1^+^ neurons. (**A**, **B**) Representative images of coronal SN sections showing GFP-expressing CRHR1 neurons co-stained with antibodies against TH and PV labelling dopaminergic and GABAergic neurons, respectively. (**C**) Representative images of neurons in the SN stained for GFP/CRHR1. Quantification of GFP/CRHR1 neurons in the SN (*p* < 0.0001, t = 6.15). Values represent mean ± SEM, * *p* < 0.05, ** *p* < 0.01, *** *p* < 0.0001. All scale bars = 200µm.

**Figure S7:**
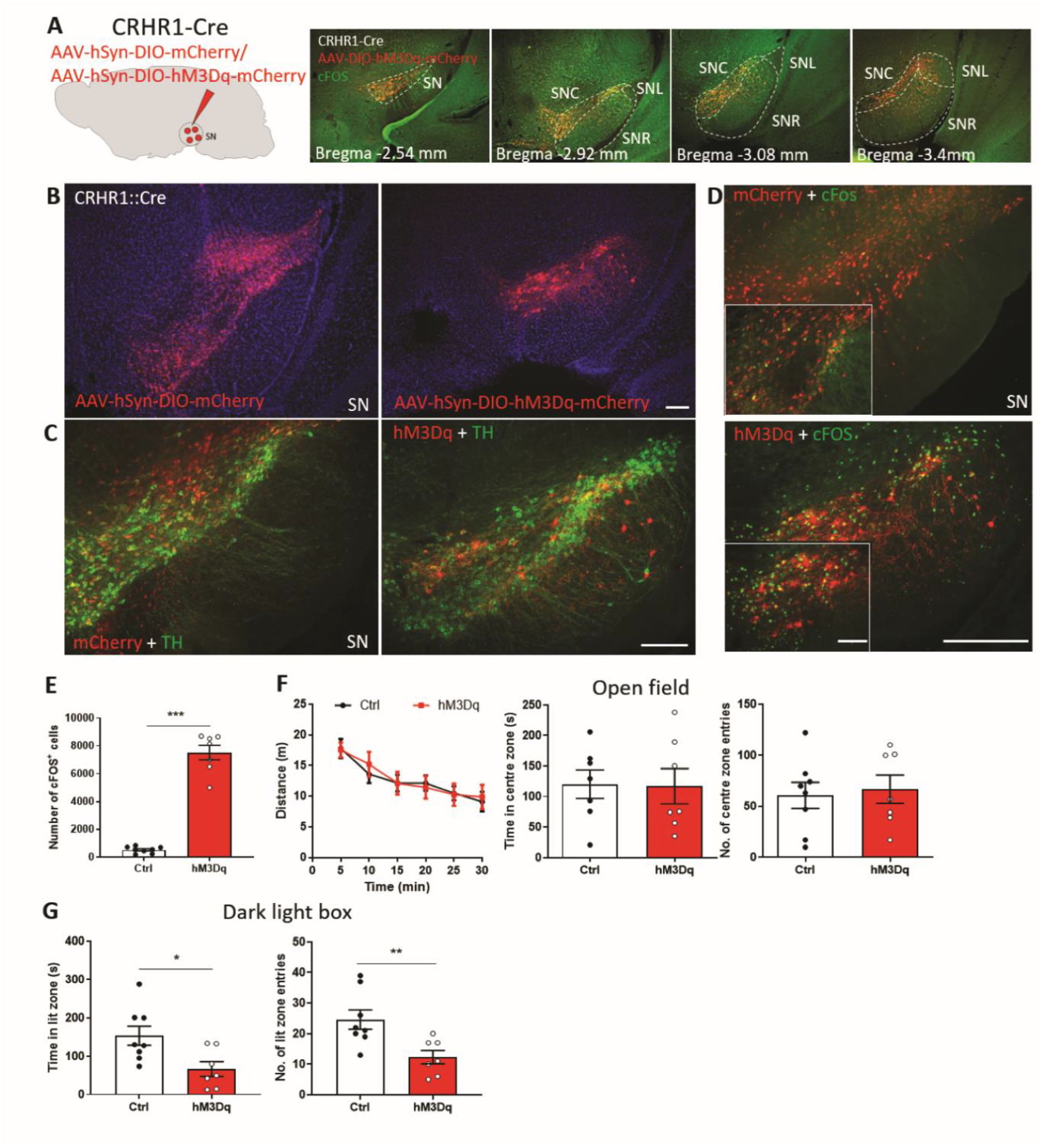
Stimulation of SN^CRHR1^ neurons does not affect arousal. (**A**) Scheme illustrating the injection of AAV_8_-hSyn-DIO-hM3Dq-mCherry or AAV_8_-hSyn-DIO-mCherry into the SN of CRHR1-Cre mice and representative images of coronal sections showing the rostro-caudal spread of mCherry expression following virus injection. (**B**) Representative images showing mCherry distribution in the SN (**C**) combined with antibody staining against TH delineating the structure of the SN. (**D**) cFOS staining in the SN of CRHR1-Cre mice expressing hM3Dq. (**E**) Quantification of cFOS^+^ cells in the SN (*p* < 0.0001, t = 11.73) 90 min following CNO administration. (**F**) Distance travelled (*F*_1,14_ = 0.01, *p* = 0.92), time spent in the centre (*p* = 0.93, t = 0.08) and entries into the centre zone (*p* = 0.74, t = 0.33) of the open field. (**G**) Time spent in lit zone (*p* = 0.018, t = 2.68) and entries into lit zone (*p* = 0.008, t = 3.09) of the dark-light box. Values represent mean ± SEM, * *p* < 0.05, ** *p* < 0.01, *** *p* < 0.0001, scale bar = 200 µm.

**Figure S8:**
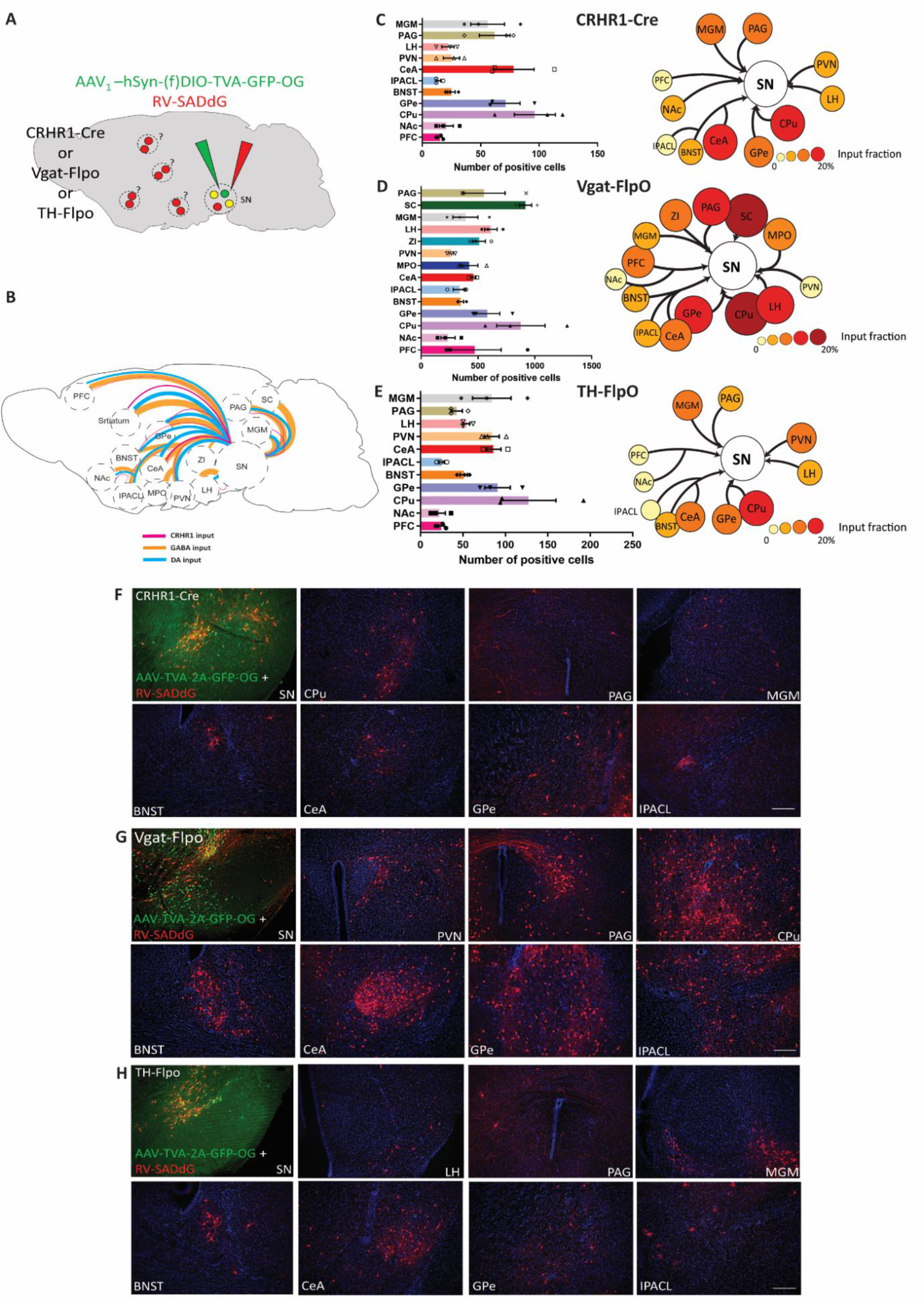
Rabies virus-mediated retrograde tracing reveals SN inputs. (**A**) Scheme depicting injection of AAV_1_-CBh-DIO-TVA-2A-GFP-OG or AAV_1_-CBh-fDIO-TVA-2A-GFP-OG and RV-SADdG into the SN of CRHR1-Cre, Vgat-FlpO or TH-FlpO mice. (**B**) Summary of rabies virus tracing results using CRHR1-Cre, Vgat-FlpO, and TH-FlpO mice illustrating inputs to the SN from different brain regions. The strength of inputs is reflected by the thickness of the lines. Quantification (**C-E)** and representative images (**F-H**) of cells labelled by rabies virus-mediated retrograde tracing in different brain regions of CRHR1-Cre, Vgat-FlpO and TH-FlpO mice. Values represent mean ± SEM, scale bar = 200 µm. Abbreviations: CPu, caudate putamen; MGM, medial geniculate thalamic nucleus; SC, superior colliculus; ZI, zona incerta.

**Figure S9:**
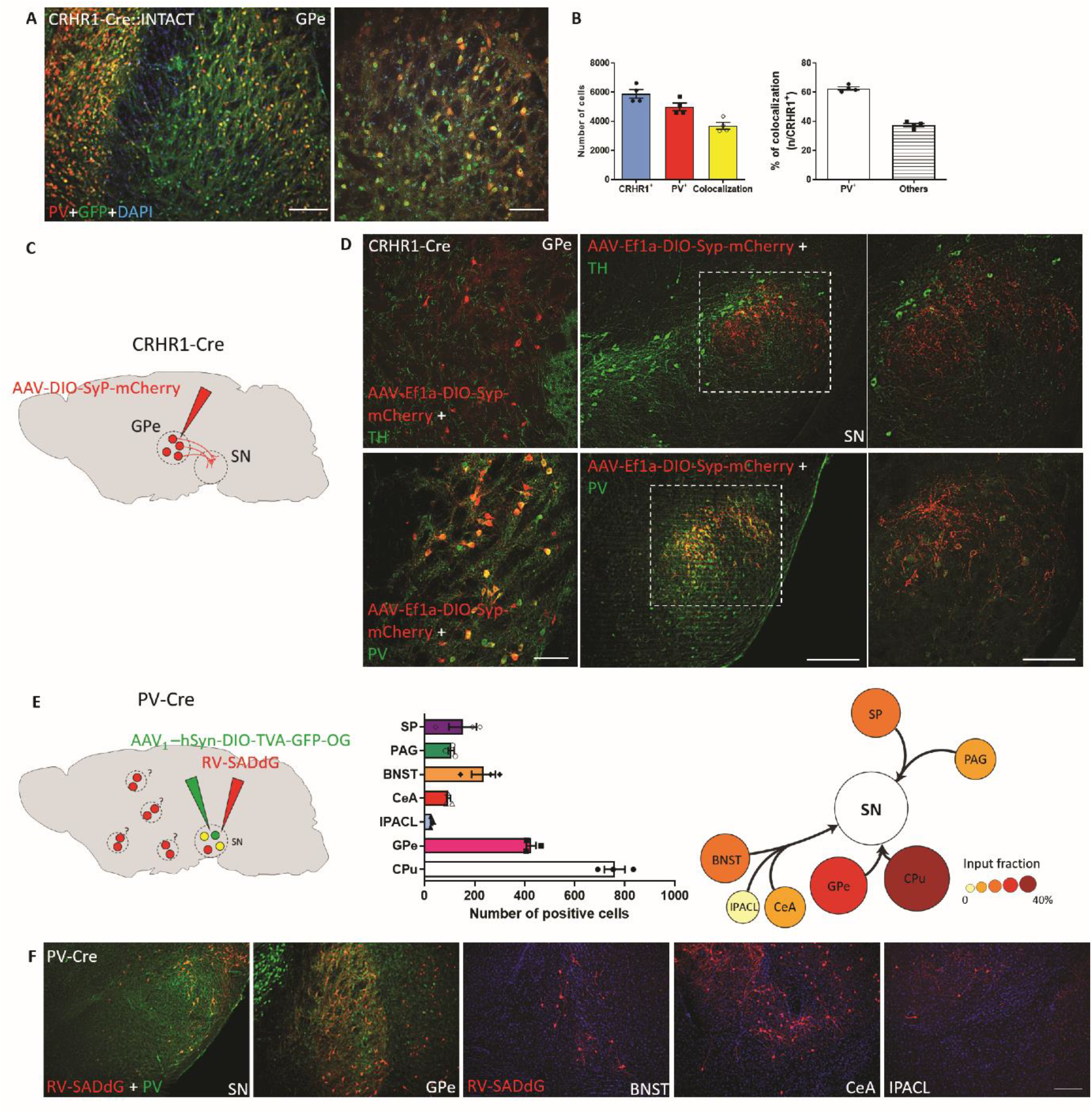
Characterization of GPe^CRHR1^ neurons. (**A**) Representative images showing nuclei of CRHR1 neurons in the GPe co-localizing with PV. (**B**) Quantification of CRHR1 and PV co-expression in the GPe. (**C**) Representative images of coronal GPe and SN section of CRHR1-Cre mice injected with AAV_8_-DIO-Syp-mCherry into the GPe of CRHR1-Cre mice stained for TH and (**D**) PV. (**E**) Representative images of coronal brain sections of the SN and retrogradely labelled brain regions. For retrograde tracing, AAV_1_-CBh-DIO-TVA-2A-GFP-OG and RV-SADdG were injected into the SN of PV-Cre mice. (**F**) Quantification of input neurons of SN^PV^ neurons throughout the brain. Values represent mean ± SEM, scale bar = 200 µm.

**Figure S10:**
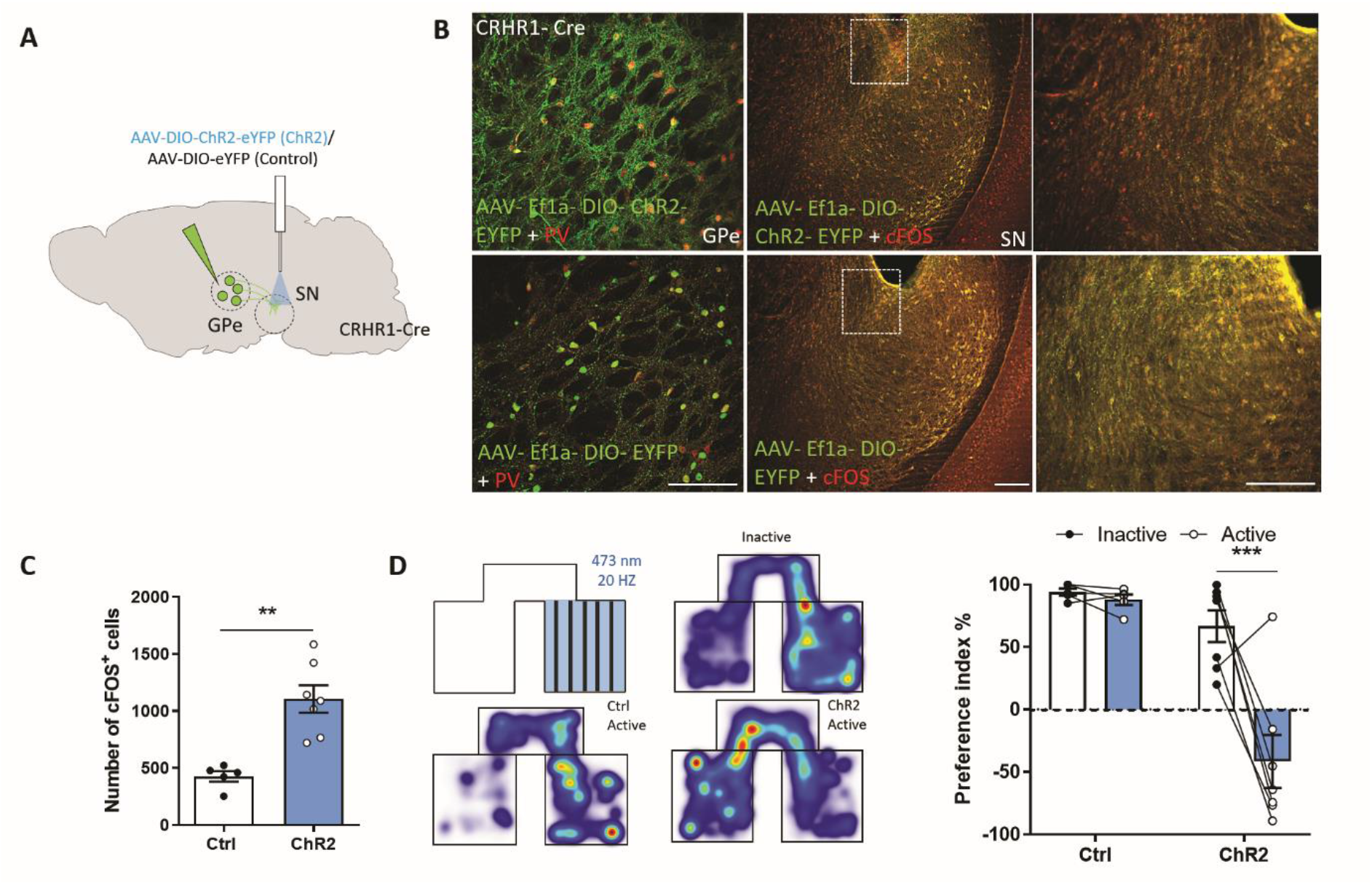
Optogenetic activation of GPe^CRHR1^ neuron terminals promotes avoidance behaviour. (**A**) Scheme illustrating injection of AAV_5_-Ef1a-DIO-ChR2-EYFP (ChR2) and AAV_5_-Ef1a-DIO-EYFP (Ctrl) into the GPe of CRHR1-Cre mice. (**B**) Representative images of coronal GPe sections from AAV injected CRHR1-Cre mice stained for PV and cFOS indicating the site of optic fibre placement in the SN. (**C**) Quantification of cFOS^+^ cells in the SN following light-induced activation of ChR2-expressing GPe^CRHR1^ neuron terminals in the SN (*p =* 0.001, t = 4.56). (**D**) RTPP paradigm with heatmaps visualizing the cumulative presence of mice in the test compartments before and after light activation. Preference index (two-way ANOVA, Bonferroni’s multiple comparison test, *p* < 0.0001, t = 5.56). Values represent mean ± SEM, * *p* < 0.05, ** *p* < 0.01, *** *p* < 0.0001. All scale bars = 200µm.

**Figure S11:**
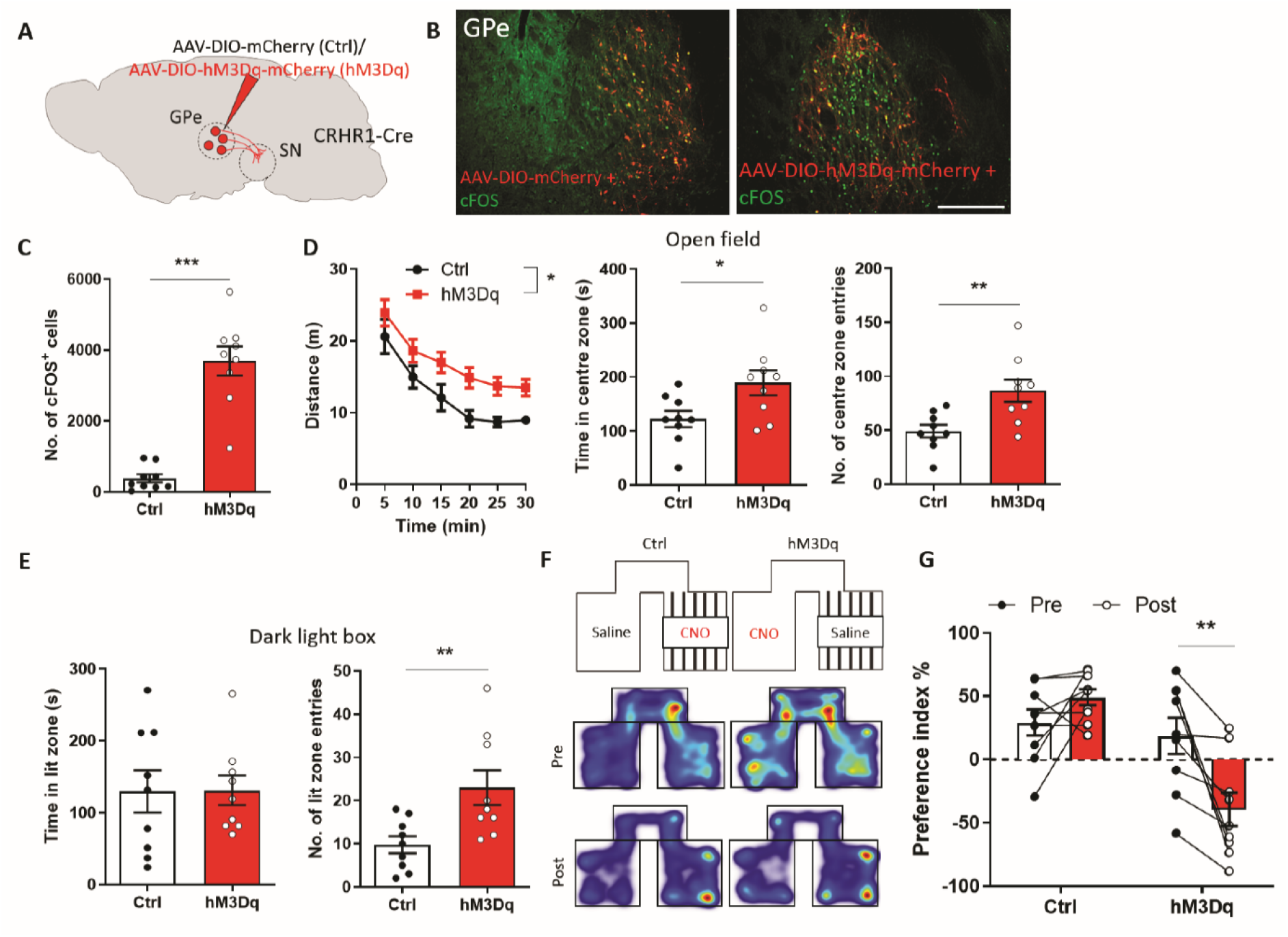
Chemogenetic activation of GPe^CRHR1^ neurons promotes arousal and avoidance behaviour. (**A)** Scheme illustrating injection of AAV_8_-hSyn-DIO-hM3Dq-mCherry (hM3Dq) or AAV_8_-hSyn-DIO-mCherry (Ctrl) into the GPe of CRHR1-Cre mice. (**B**) Representative coronal GPe sections stained for cFOS and (**C**), quantification of cFOS^+^ cells 90 min following CNO administration (*p* < 0.0001, t = 7.81). (**D-G**) CNO treatment induces arousal, delayed habituation and avoidance behaviour in hM3Dq-expressing animals. (**D**) Distance travelled (n = 9, repeated measures two-way ANOVA, *F*_1,14_ = 6.54, *p* = 0.022), time spent in the centre zone (*p* = 0.027, t = 2.43) and entries into the centre zone (*p* = 0.006, t = 3.15) of the OF. (**E**) Time spent in the lit zone (*p* = 0.95, t = 0.04) and entries into the lit zone (*p* = 0.009, t = 2.94) of the DaLi. (**F**) CPP paradigm with heatmaps visualizing the cumulative presence of mice in the test compartments pre and post CNO treatment. (**G**) Preference index (two-way ANOVA, Bonferroni’s multiple comparison test, *p* = 0.006, t = 3.59). Values represent mean ± SEM, * *p* < 0.05, ** *p* < 0.01, *** *p* < 0.0001. All scale bars = 200µm.

**Figure S12:**
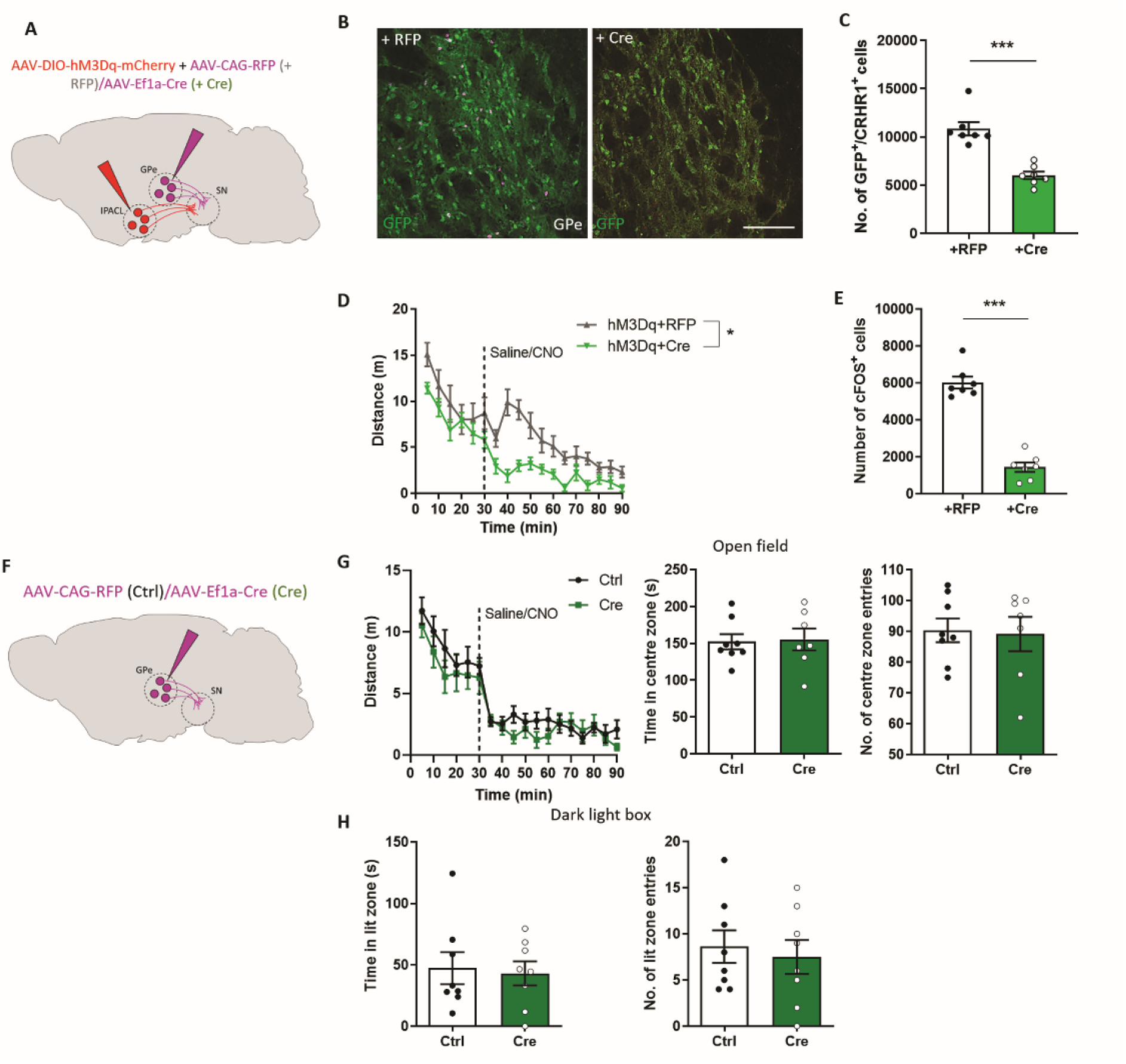
CRHR1 in GPe^CRHR1^ neurons contributes to arousal and avoidance behaviour. (**A**) Scheme illustrating injection of AAV_8_-hSyn-DIO-hM3Dq-mCherry into the IPACL and of AAV_1_-Ef1a-Cre (hM3Dq + Cre) or AAV_1_-Ef1a-RFP (hM3Dq + Ctrl) into the GPe of CRH-Cre::CRHR1^N-EGFP^ mice. (**B**) Representative coronal images and (**C**) quantification of GFP/CRHR1 neurons in the GPe (*p* < 0.0001, t = 6.15). (**D**) Locomotor activity of CRHR1-Cre mice injected with AAV_8_-hSyn-DIO-hM3Dq (hM3Dq) or AAV_8_-hSyn-DIO-mCherry (Ctrl) into the GPe in a 90-min open field test (repeated measures two-way ANOVA, *F*_1,16_ = 7.4, *p* = 0.014). (**E**) Quantification of cFOS^+^ cells in the GPe of hM3Dq mice 90 min following CNO administration (*p* < 0.0001, t = 11.2). (**F**) Schematic illustration of control experiment showing injection of AAV_1_-Ef1a-Cre (Cre) or AAV_1_-Ef1a-RFP (Ctrl) into the GPe of CRH-Cre::CRHR1^N-EGFP^ mice. (**G**) Distance travelled (*F*_1,14_ = 0.31, *p* = 0.58), time spent in centre (*p* = 0.86, t = 0.18) and entries into the centre zone (*p* = 0.86, t = 0.19) of the open field. (**H**) Time spent in lit zone (*p* = 0.8, t = 0.25) and entries into lit zone (*p* = 0.67, t = 0.44) of the dark-light box. Values represent mean ± SEM, * *p* < 0.05, ** *p* < 0.01, *** *p* < 0.0001, scale bar = 200 µm.

**Figure S13:**
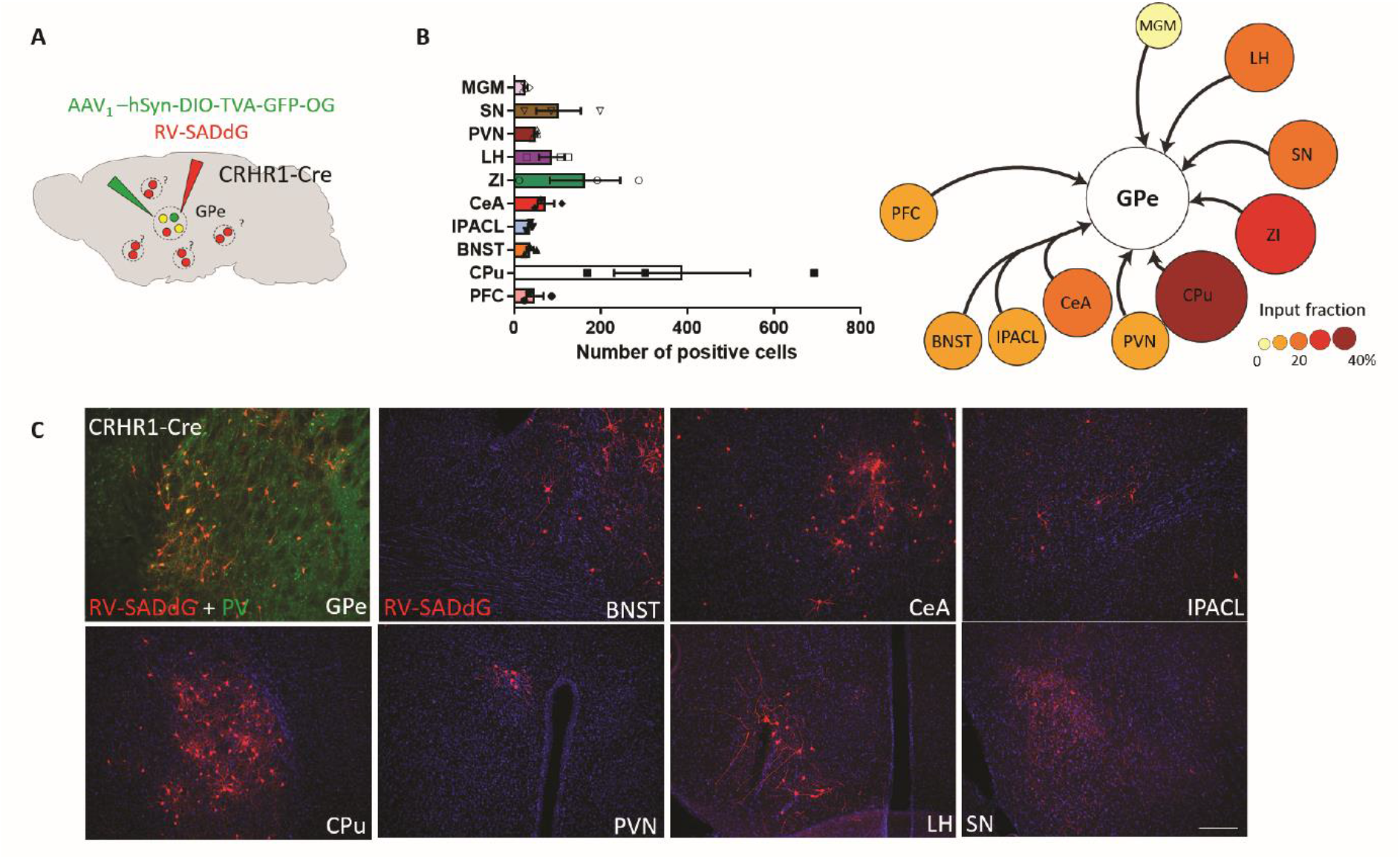
Rabies virus-mediated retrograde tracing reveals inputs of GPe^CRHR1^ neurons. (**A**) Scheme depicting injection of AAV_1_-CBh-DIO-TVA-2A-GFP-OG and RV-SADdG into the GPe of CRHR1-Cre mice. Representative images of coronal brain sections of the GPe and retrogradely labelled brain regions. For retrograde tracing, AAV_1_-CBh-DIO-TVA-2A-GFP-OG and RV-SADdG were injected into the SN of CRHR1-Cre mice. (**B**) Quantification of input neurons of GPe^CRHR1^ neurons throughout the brain. (**C**) Representative images of cells labelled by rabies virus-mediated retrograde tracing in different brain regions of CRHR1-Cre mice. Values represent mean ± SEM. All scale bars = 200µm.

**Figure S14:**
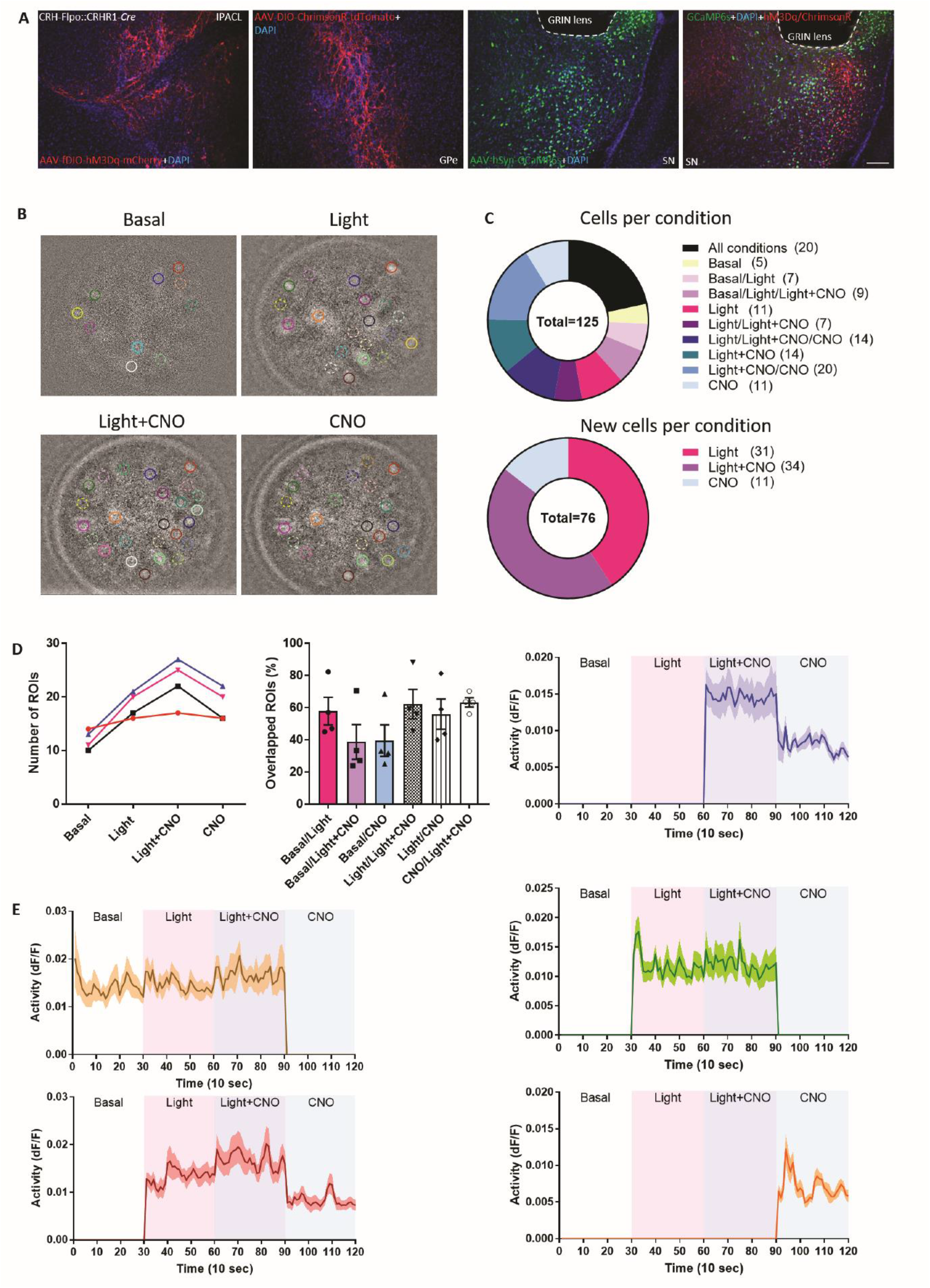
In vivo calcium imaging reveals differential cellular responses depending on stimulation conditions. **(A)** Representative images showing sites of virus injection and GRIN lens implantation in the lateral SN. (**B**) Representative images showing ROIs detected under different conditions during miniscope recording. (**C**) Pie charts showing the distribution of cells and number of newly appeared cells during different conditions. (**D**) Number of ROIs captured from each animal under different stimulation conditions. Number of overlapping ROIs under different stimulation conditions. (**E**) Traces of clusters of cells activated during different stimulation conditions. Values represent mean ± SEM, scale bar = 200 µm.

## Notes

### Competing Interest Statement

The authors have declared no competing interest.

